# Mapping and explaining wolf recolonization in France using dynamic occupancy models and opportunistic data

**DOI:** 10.1101/099424

**Authors:** Julie Louvrier, Christophe Duchamp, Eric Marboutin, Sarah Cubaynes, Rémi Choquet, Christian Miquel, Olivier Gimenez

## Abstract

While large carnivores are recovering in Europe, assessing their distributions can help to predict and mitigate conflicts with human activities. Because they are highly mobile, elusive and live at very low density, modeling their distributions presents several challenges due to i) their imperfect detectability, ii) their dynamic ranges over time and iii) their monitoring at large scales consisting mainly of opportunistic data without a formal measure of the sampling effort. Not accounting for these issues can lead to flawed inference about the distribution.

Here, we focused on the wolf (*Canis lupus*) that has been recolonizing France since the early 90’s. We evaluated the sampling effort *a posteriori* as the number of observers present per year in a cell based on their location and professional activities. We then assessed wolf range dynamics from 1993 to 2014, while accounting for species imperfect detection and time- and space-varying sampling effort using dynamic site-occupancy models.

Ignoring the effect of sampling effort on species detectability led to underestimating the number of occupied sites by 50% on average. Colonization increased with increasing number of occupied sites at short and long-distances, as well as with increasing forest cover, farmland cover and mean altitude. Colonization decreased when high-altitude increased. The growth rate, defined as the number of sites newly occupied in a given year divided by the number of occupied sites in the previous year, decreased over time, from over 100% in 1994 to 5% in 2014. This suggests that wolves are expanding in France but at a rate that is slowing down. Our work shows that opportunistic data can be analyzed with species distribution models that control for imperfect detection, pending a quantification of sampling effort. Our approach has the potential for being used by decision-makers to target sites where large carnivores are likely to occur and mitigate conflicts.

## Introduction

Large carnivores are often considered as key elements for maintaining ecosystems. Because of their high position in the trophic chain, their extinction can lead to trophic cascades and detrimental changes in species abundance and functioning of ecosystems (Ripple et al. 2014). Once widespread in Europe, many populations of large carnivores were extirpated over the last century, mainly due to interferences with human activities (Breitenmoser 1998, Ripple et al. 2014). Since the 1970s, all large carnivores have recovered, benefiting from legal protection and the recovery of wild ungulate populations, resulting in most of the European countries hosting at least one viable population of large predators (Chapron et al. 2014). Often used as a conservation success story, the recovery of large carnivores in human-dominated areas comes with challenges, including the question of whether there are any sufficiently large and functional areas left for viable populations (Packer et al. 2013). Another issue is how to coordinate management of these species at large scales, possibly across borders (Linnell and Boitani 2012, Bischof et al. 2015), in particular in the context of international treaties and directives (e.g. the Habitats Fauna Flora European Directive).

In this context, mapping the distribution of a species can help to predict and mitigate conflicts. Species distribution models (SDMs) have become important tools in the ecological, biogeographical and conservation fields (Guisan and Thuiller 2005). By correlating presence-only or presence-absence data of a species to environmental factors, SDMs provide an understanding of habitat preferences and predictions on future species distribution. This is especially relevant for species involved in conflicts, since predicting their future presence can help targeting contentious areas and guide management to reduce conflicts (Guillera-Arroita et al. 2015). However, the monitoring of large carnivores remains challenging to carry out in the field because these species live at low density and occupy wide areas (Gitlleman et al. 2001). Therefore, assessing the distribution of these species comes with methodological challenges.

First, standard SDMs such as Maxent (Phillips et al. 2006) rely on the assumption of perfect detection, assuming that the focal species is detected everywhere it is present (Yackulic et al. 2013). Going undetected at a given site does not necessarily mean that this species is absent from that site, but rather that it may simply be missed for various reasons related to observer abilities, habitat characteristics or species level of activity (Kéry et al. 2010, Kéry 2011). Ignoring the issue of imperfect detection can result in false absences that lead to flawed inference in two ways: i) the distribution maps are biased by underestimating actual presences and can misrepresent certain viable habitat features that are falsely identified as unfavorable (Kéry and Schaub 2011, Lahoz-Monfort et al. 2014); ii) there may be confusion in identifying the drivers of the species distribution when detection depends on environmental explanatory variables that are independent from the variables influencing the species’ actual presence. For instance, if altitude has a negative effect on detection but not on presence, then as altitude gets higher, the species is less likely to be detected which can lead to erroneous conclusions that the species prefers lower altitudes (Lahoz-Monfort et al. 2014). To cope with this first issue, single-season or static site-occupancy models were developed (Mackenzie et al. 2006) and have been widely used for carnivores (e.g., Thorn et al. 2011 for brown hyenas *Hyaena brunnea*; Long et al. 2010 for black bears *Ursus americanus*, fishers *Martes pennant* and bobcats *Lynx rufus*; Sunarto et al. 2012 for Sumatran Tigers *Panthera tigris sumatrae*). Based on spatial and temporal replicated sampling of the target species, these models allow assessing the effects of environmental factors on species occupancy, while making the distinction between non-detections and true absences via the estimation of species detectability.

Second, most SDMs are implicitly based on the ecological niche concept (Grinnell 1917; Hutchinson 1957) and therefore rely on two main hypotheses: i) the species is present in areas where environmental conditions are the most favorable and ii) dispersal is not a limiting factor (Jeschke and Strayer 2006). However, expanding species are often absent from an area not because conditions are unfavorable but because they have not yet dispersed to this area, or because of geographical barriers or dispersal constraints (Araújo and Guisan 2006). Hence, static SDMs ignore important dynamic processes, which may lead to bias in the resulting distributions (Yackulic et al. 2015). Static SDMs should therefore not be used for predictions (Zurell et al. 2009). To deal with this second issue, occupancy models have been extended (Mackenzie et al. 2003, Royle and Kéry 2007) to account for the influence of dynamic processes such as colonization and extinction on the species range dynamics (Mackenzie et al. 2003). So-called multi-season or dynamic site-occupancy models are increasingly used to assess the range dynamics of expanding or invasive species (e.g., Bled et al. 2011 for the hadeda ibis *Bostrychia hagedash* and Broms et al. 2016a for the common myna *Acridotheres tristis*), but remain rarely applied to carnivores (e.g., Marcelli and Fusillo 2012 for the Eurasian otter *Lutra lutra* or Miller et al. 2013 for the grey wolves *Canis lupus*).

Third, data collection is particularly costly if not unfeasible for elusive species that need wide areas due to the large presence area required for sampling. In this context, citizen science is considered as an efficient source of information to assess changes in a species distribution by covering wide areas (Schmeller et al. 2009). However, data from citizen science are often collected with protocols that do not control for variation in the sampling effort i) in time: a site can be sampled by several observers during a given year and not the following year and ii) in space: given two sites where the species is present, if the sampling effort is higher in one site, this might lead to recording a false absence in the site with lower sampling effort (Kéry et al. 2010). As a consequence, if sampling effort is not controlled for, detectability can be underestimated as well as its variability, leading to an overestimation of the distribution area (Van Strien et al. 2013).

Static and dynamic occupancy models hold promise to analyze population trends from opportunistic data because the data collection process is formally incorporated (Isaac et al. 2014). Occupancy models are divided into two types of process; i) the ecological one, governing dynamic occupancy processes (initial occupancy, local extinction and local colonization) and ii) the observation process, governing the data collection process (Mackenzie et al. 2003). However, to address the third issue and apply occupancy models to opportunistic data, one needs to differentiate between a site that was not sampled and a site that was sampled but the species was not detected. In the case of several species being monitored, the detection of a species in a site informs about the non-detection of other species because this site is known to have been sampled (Van Strien et al. 2013). This no longer holds for single-species settings, and the assumption is sometimes made that all sites where at least one detection occurred are sampled throughout the whole duration of the study (Molinari-Jobin et al. 2012, Rich et al. 2013).

Here, we considered Grey wolves (*Canis lupus*) as a case study to illustrate the challenges in using opportunistic data and SDMs to infer the range dynamics of large carnivores. Wolves disappeared in most of the Western European countries during the twentieth century (Promberger and Schroder 1993; Boitani 2010) except in Spain, Portugal and Italy (Boitani and Cucci 1993). The species naturally recolonized the French Alps from the remaining Italian population (Valière et al. 2003, Fabbri et al. 2007). Because the species is protected by law while being a source of conflicts with sheepherding, its recolonization process needs to be carefully monitored.

Our main objective was to describe and determine the drivers of wolves recolonization pattern in France between 1993 and 2014. To account for imperfect detection, we built a dynamic site-occupancy model (Mackenzie et al. 2006) and analyzed opportunistic data collected by a network of trained volunteers since 1992. To do so, we built *a posteriori* the sampling effort to account for biases in data collected through citizen science. To describe the recolonization process over time, we addressed two main questions: (i) What are the environmental and biological factors influencing colonization and extinction probabilities? (ii) How can sampling effort be inferred *a posteriori*, i.e. after the data were collected, and to what extent does sampling effort correlate with detection probability?

## Methods

### Study species and area

The first wolf (*Canis lupus*) occurrence was detected in France in the early 1990s as a consequence of the Italian population’s expansion (Valière et al. 2003, Ciucci et al. 2009). The species then spread outside the Alpine mountains to reach the Pyrenees and the Massif Central westward first in 1999, and the Vosges Mountains northward from 2011. The wolf is an opportunist species that can adapt its diet depending on available prey species (Poulle et al. 1997, Imbert el al. 2016). In areas with livestock farming, strong interactions between wolf presence and sheep breeding usually occur. The study area mostly covered Eastern France and a major part of Central France (Fig. 1).

**Figure 1:**
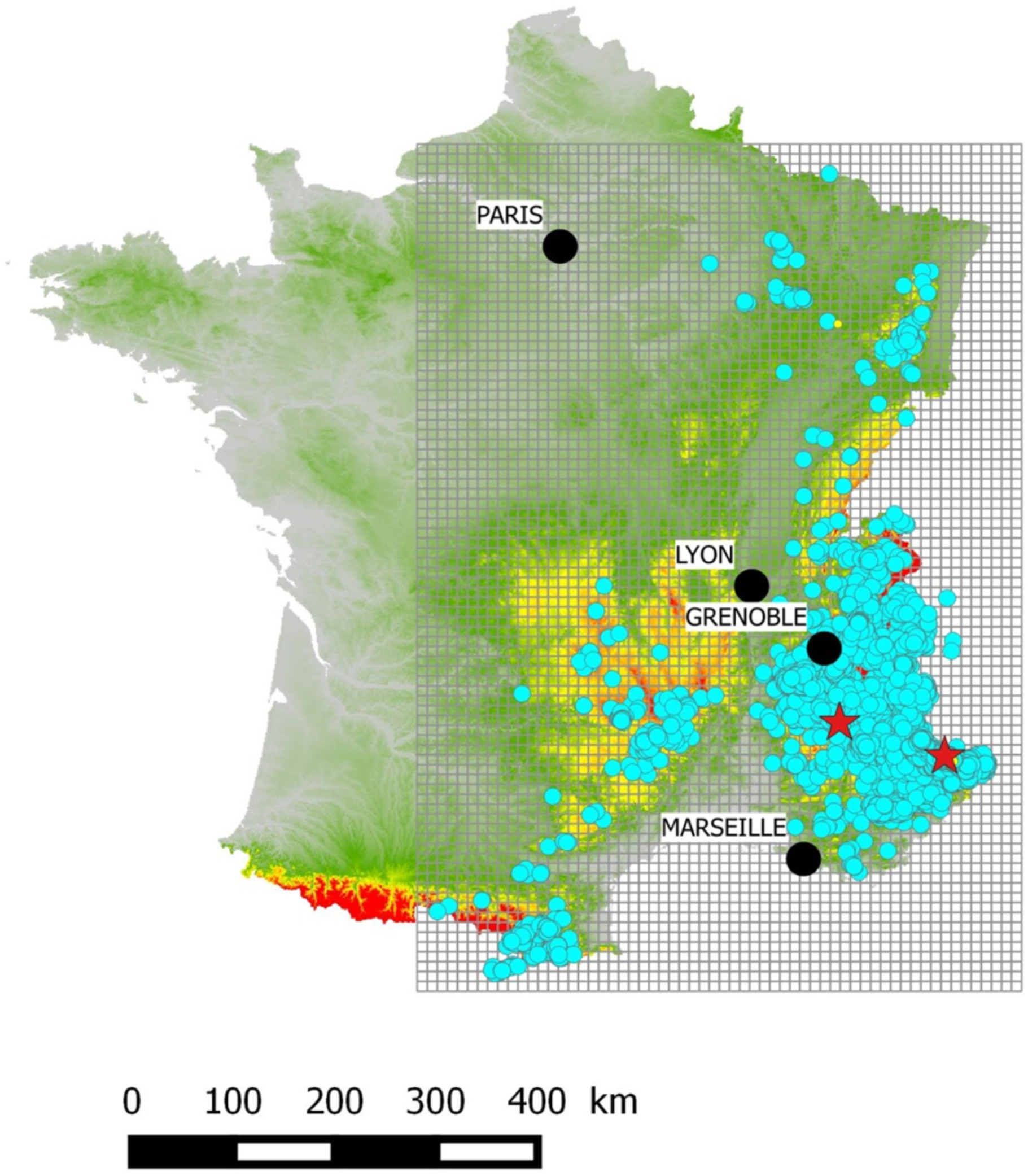
Maps of cumulated species detections (blue dots) for the period 1993-2014. Red stars represent the first detections made in 1992. Sites were defined as 10x10km cells within a grid covering all detections. Red areas represent mountainous areas with an altitude higher than 1500 meters.

### Data collection

Wolf detection data were made of presence signs sampled all year long from 1992 to 2014 thanks to a network of professional and non-professional observers. The network size has increased from a few hundred people in 1994, up to 3000 wolf experts in 2015. Every observer is trained during a 3-day teaching course led by the French National Game and Wildlife Agency (ONCFS) to document signs of the species presence (Duchamp et al. 2012). Presence signs went through a standardized control process combining genetic identification tools, and validation standards to prevent misidentification (Duchamp et al. 2012). For every presence sign, the date and location of collection were stored in a geo-referenced database. These data are considered opportunistic in the sense that monitoring occurs all year long in an extensive manner without explicitly quantifying the sampling effort.

### Dynamic site occupancy models

To model the colonization dynamics of wolf, we used dynamic site-occupancy models (Mackenzie et al. 2003). These models allow the quantification of species occupancy while correcting for imperfect species detectability based on repeated sampling in time and space. We defined sampling units as 10x10km cells, which appears to be the best option in the context of our study (Marboutin et al. 2010) and also is the recommended surface to produce maps of presence by the European Union (E. C. 2006). Site occupancy models rely on several assumptions, including the closure assumption which states that the ecological state of a site (whether it is occupied or not) remains unchanged through occasions (or surveys) *j* within a year *k*. During year *k*, sites were monitored mainly in winter from December to March, the most favorable period to detect the species between the two peaks of dispersal events in spring and fall (Mech and Boitani 2010). We defined the secondary occasions *j* as December, January, February and March and *y_i,j,k_*, the observed state of site *i* equal to 1 if at least one sign of presence was found at site *i* during occasion *j* in the year *k* (and 0 otherwise).

We considered a state-space formulation of the dynamic occupancy model (Royle and Kéry 2007) in which the model is viewed as the combination of (i) the ecological process that involves the latent ecological state of a site, i.e. whether it is occupied or not; (ii) the observation process that leads to the detections or non-detections by the observer conditional on the state of the system. The colonization probability *γ_i,k_* is the probability that an empty site *i* during year *k* becomes occupied during year *k*+1, while the extinction probability *ε_i,k_* is the probability that an occupied site *i* during year *k* becomes empty during year *k*+1. We define *z_i_*,_1_ as the initial latent state of site *i* as being drawn from a Bernoulli distribution with the success probability being *Ψ_i_*,_1_, *z_i_*,_1_ ~ *Bernoulli* (*Ψ_i_*,_1_). All other latent states *z_i,k_* for *k* > 1 are drawn from a Bernoulli distribution as *z_i,k_*_+1_ | *z_i,k_* ~ *Bernoulli* (*z_i,k_* (1 - *ε_i,k_*) + (1 - *z_i,k_*) *γ_i,k_*). On top of the ecological process stands the observation process, in which the detections/non-detections are drawn from a Bernoulli distribution *y_i,j,k_*\*z_i,k_* ~ *Bernoulli*(*z_i,k_ p_i,j,k_*) where *p_i,j,k_* is the probability that the species is detected at site *i* for an occasion *j* during year *k*. The state-space formulation is appealing as it makes explicit the latent states *z_i,k_* that can be used to build distribution maps.

### Sampling effort

Monitoring the range expansion of wolves at the country level prevented us from implementing any standardized experimental sampling design. Instead, the presence signs were sampled in an opportunistic way and the sites were defined *a posteriori*. When dealing with detection-only data, various approaches have been adopted to infer the non-detections. In the context of species-list protocols, if other species are detected at a site but not the focal species, one can assume that observers were present but did not detect the species of interest and, hence a non-detection for this species is recorded at this given site (Kéry et al. 2010). In the context of single-species monitoring, several authors have assumed that the observation effort was sufficient enough to make the assumption that all occupied cells were monitored during the study period in any year (e.g., Molinari-Jobin et al. 2012). We adopted an original approach to infer the non-detections based on the available qualitative information on the observers. When entering the network, observers attended a 3-day training session to learn how to identify the species and how it is monitored (Duchamp et al. 2012). During these training sessions, we recorded the observers’ personal and professional address, socio-professional category and entry date into the network. The entry date was used to quantify how many observers were present in the network each year. We calculated a circular buffer for the prospection area for each observer based on a radius specific to his/her socio-professional category and a center located at his/her address (Supplementary material, table A1). For instance, for an observer belonging to the category 1 (departmental authority) whose address was located in the French Department number 39, his/her prospection area would be 4 999 km^2^, which is the size of the Department (Supplementary material, Fig. A1 and Table A2). For this observer, a circular buffer was built with a radius calculated as
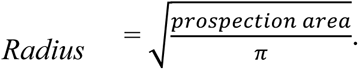

**Table A1:** Size of prospection areas as a function of socio-professional category of observers. Observers were classified according to 8 entities to capture the diversity of their professional and personal field activities. People working for the departmental authorities (Category 1) display a field effort all over that departmental area. Observers belonging to the category 2 are state employees affected to the protected area they are working in. Details were not given for Regional Natural Park agents and Natural reserve agents. Their prospection area corresponds to the mean area of the protected area they are affiliated to. ONCFS agents (category 8) are attributed half a French Department as field areas when assigned for species monitoring. ONF agents (category 9) are attributed 1/10 of a French Department. Farmers (category 4) and hunters (category 5) usually focus on the restricted area (“municipality”) where they farm, breed sheep or hunt. Scientists (category 3), members of a naturalist association (category 6) and volunteers (category 7) were given ¼ of their affiliated department as their main activity might not be focused on species monitoring.

| Socio-professional category | Prospection area | Number of observers |
| --- | --- | --- |
| Administrative (e.g., department authorities) **(category 1)** | Area of the affiliated French department | 230 |
| National Park agent **(category 2)** | Area of the affiliated National Park | 156 |
| Regional Natural Park agent **(category 2)** | 450 km<sup>2</sup> | 69 |
| Natural Reserve agent **(category 2)** | 10 km<sup>2</sup> | 20 |
| Scientist **(category 3)** | $\frac{1}{4}$ of the affiliated department's surface | 6 |
| Agricultural profession **(category 4)** | Area of the affiliated municipality | 59 |
| Hunter **(category 5)** | Area of the affiliated municipality | 313 |
| Member of a naturalist association **(category 6)** | $\frac{1}{4}$ of the affiliated department's surface | 216 |
| Volunteer **(category 7)** | $\frac{1}{4}$ of the affiliated department's surface | 311 |
| ONCFS agent **(category 8)** | $\frac{1}{2}$ of the affiliated department's surface | 513 |
| French Forest Agency (ONF) agent **(category 9)** | 1/10 of the affiliated department's surface | 284 |

**Table A2:** French departments where observers are present in 2014 along with their area and the number of observers affiliated to each department depending on their socio-professional category. Categories 2, 4 and 5 are not shown because their prospection areas do not depend on the size of the affiliated department. See also Figure A1.

| Department code | Surface (km<sup>2</sup>) | Category 1 | Category 3 | Category 6 | Category 7 | Category 8 | Category 9 |
| --- | --- | --- | --- | --- | --- | --- | --- |
| 1 | 5762 | 5 | 0 | 8 | 8 | 22 | 11 |
| 4 | 6925 | 20 | 0 | 5 | 24 | 12 | 12 |
| 5 | 5549 | 8 | 0 | 4 | 17 | 19 | 6 |
| 6 | 4299 | 5 | 0 | 0 | 18 | 14 | 1 |
| 7 | 5529 | 5 | 0 | 6 | 0 | 12 | 4 |
| 8 | 5229 | 0 | 0 | 0 | 3 | 1 | 0 |
| 9 | 4890 | 0 | 0 | 4 | 1 | 11 | 7 |
| 10 | 6004 | 0 | 1 | 1 | 0 | 2 | 0 |
| 11 | 6139 | 0 | 0 | 0 | 0 | 3 | 2 |
| 12 | 8735 | 4 | 0 | 0 | 3 | 11 | 4 |
| 13 | 5087 | 0 | 0 | 1 | 1 | 10 | 1 |
| 15 | 5726 | 5 | 0 | 4 | 3 | 11 | 2 |
| 19 | 5857 | 0 | 0 | 0 | 0 | 2 | 0 |
| 21 | 8763 | 2 | 0 | 0 | 1 | 11 | 0 |
| 23 | 5565 | 0 | 0 | 0 | 0 | 1 | 0 |
| 24 | 9060 | 0 | 0 | 0 | 0 | 2 | 0 |
| 25 | 5234 | 14 | 0 | 11 | 4 | 22 | 11 |
| 26 | 6530 | 9 | 0 | 6 | 10 | 14 | 18 |
| 30 | 5853 | 4 | 0 | 3 | 0 | 7 | 1 |
| 31 | 6309 | 1 | 1 | 0 | 6 | 12 | 1 |
| 32 | 6257 | 0 | 0 | 0 | 1 | 8 | 0 |
| 34 | 6101 | 0 | 1 | 1 | 0 | 3 | 0 |
| 38 | 7431 | 6 | 1 | 21 | 29 | 28 | 23 |
| 39 | 4999 | 5 | 0 | 16 | 7 | 12 | 14 |
| 42 | 4781 | 8 | 0 | 1 | 4 | 3 | 3 |
| 43 | 4977 | 2 | 0 | 0 | 1 | 4 | 0 |
| 46 | 5217 | 0 | 0 | 0 | 0 | 1 | 0 |
| 47 | 5361 | 0 | 0 | 0 | 0 | 3 | 0 |
| 48 | 5167 | 17 | 0 | 5 | 6 | 14 | 8 |
| 54 | 5246 | 5 | 0 | 9 | 3 | 7 | 4 |
| 55 | 6211 | 1 | 0 | 1 | 1 | 23 | 3 |
| 57 | 6216 | 4 | 0 | 4 | 6 | 17 | 1 |
| 58 | 6817 | 0 | 0 | 1 | 0 | 0 | 0 |
| 59 | 5743 | 0 | 0 | 0 | 0 | 0 | 0 |
| 63 | 7970 | 0 | 1 | 5 | 1 | 6 | 1 |
| 65 | 4464 | 0 | 0 | 0 | 0 | 2 | 0 |
| 66 | 4116 | 6 | 0 | 7 | 18 | 10 | 11 |
| 67 | 4755 | 11 | 0 | 19 | 16 | 31 | 14 |
| 68 | 3525 | 28 | 0 | 20 | 44 | 16 | 17 |
| 69 | 3249 | 1 | 0 | 1 | 2 | 4 | 0 |
| 70 | 5360 | 10 | 0 | 11 | 1 | 12 | 12 |
| 71 | 8575 | 0 | 0 | 1 | 2 | 7 | 1 |
| 73 | 6028 | 7 | 0 | 5 | 17 | 17 | 33 |
| 74 | 4388 | 6 | 0 | 16 | 29 | 20 | 13 |
| 81 | 5758 | 0 | 0 | 0 | 0 | 4 | 1 |
| 82 | 3718 | 3 | 0 | 0 | 0 | 3 | 0 |
| 83 | 5973 | 3 | 1 | 2 | 3 | 15 | 1 |
| 84 | 3567 | 2 | 0 | 1 | 0 | 10 | 5 |
| 87 | 5520 | 0 | 0 | 0 | 1 | 0 | 0 |
| 88 | 5874 | 16 | 0 | 12 | 17 | 25 | 33 |
| 89 | 7427 | 0 | 0 | 1 | 0 | 0 | 0 |
| 90 | 609 | 7 | 0 | 3 | 3 | 9 | 5 |

**Figure A1:**
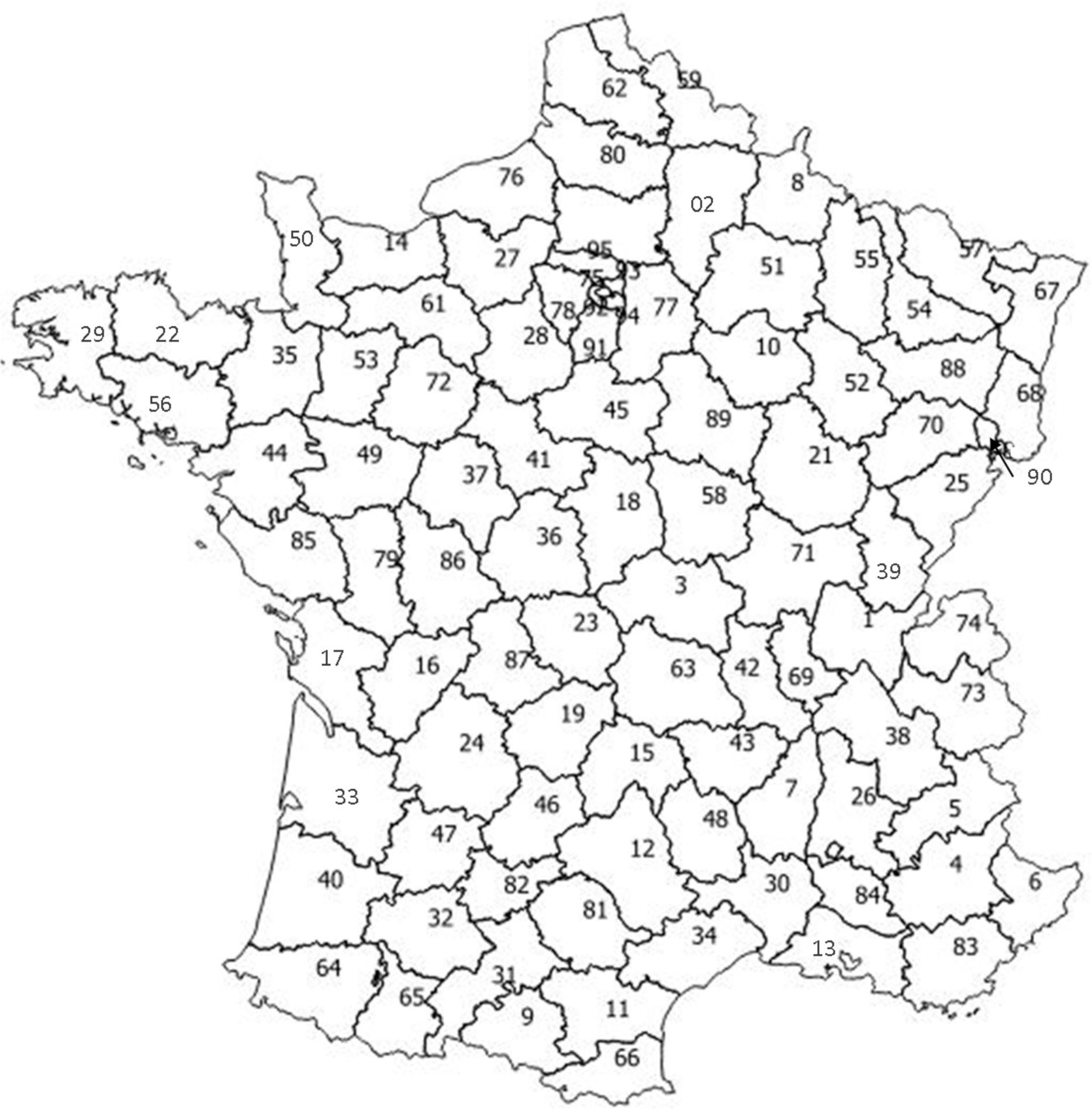
Map of French departments with the identity code used in Table A2.

For each 10x10km cell, we then calculated the number of observers monitoring the species per year by summing the number of prospection areas overlapping the cell (supplementary material, Fig. A2). We set the species detection probability at a site to zero when the sampling effort was null in that site, i.e. no observers were present in that cell. When at least one observer was found in a cell in a given year, we considered that sampling occurred, hence concluding that a presence sign found at a particular occasion this year was a detection, and a non-detection otherwise. We expected that with more observers per site per year, the species was more likely to be detected, in other words that the sampling effort had a positive effect on the detection parameter. We performed a sensitivity analysis to assess how a change in the construction of the sampling effort influenced the model parameter estimates (Supplementary material, Fig. A3).

**Figure A2:**
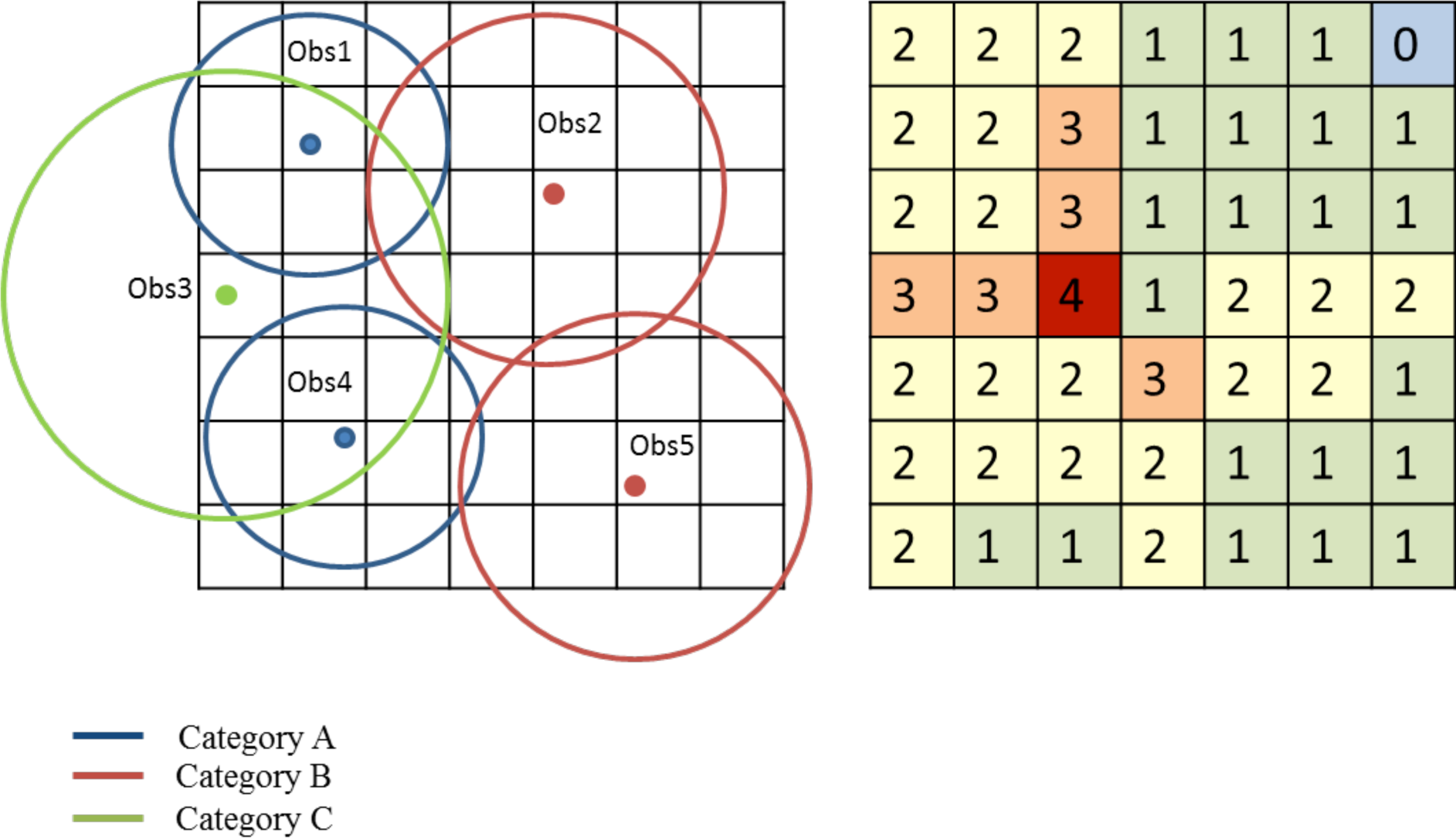
Schematic representation of how the sampling effort was calculated. Left: Observers were plotted according to their address. A circular buffer was affiliated to each observer with a surface equal to the prospection area following Tables A1 and A2. Right: resulting sampling effort calculated as the sum of observers sampling in each cell.

**Figure A3:**
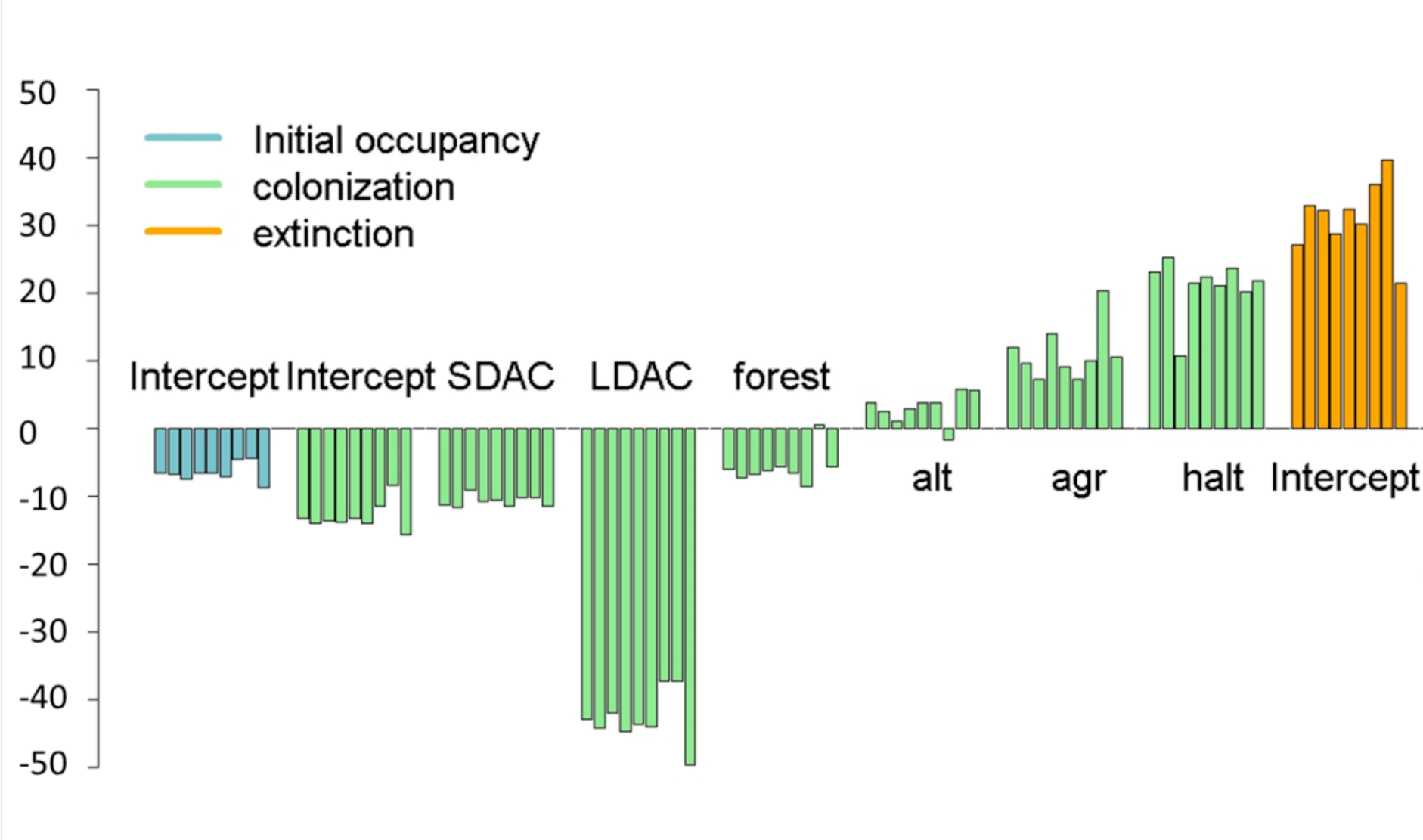
Analysis of sensitivity to the sampling effort definition. Each bar within a barplot represents the percentage of change (in %) in parameter estimates when compared to our original definition of the sampling effort after modification of the prospection area for one category of observers, in the following order: category 6: 1/10 of the affiliated department’s surface; category 6: 1/2 of the affiliated department’s surface; category 6: 100% of the affiliated department’s surface; category 7: 1/10 of the affiliated department’s surface; category 7: 1/2 of the affiliated department’s surface; category 7: 100% of the affiliated department’s surface; category 8: 1/10 of the affiliated department’s surface; category 8: 1/4 of the affiliated department’s surface; category 8: 100% of the affiliated department’s surface. The model best supported by the data is used throughout these analyses.

### Habitat covariates

Wolves can adapt to a large range of different habitats, which makes it difficult to identify specific factors that may influence the species’ presence at a site (Mech and Boitani 2010). However, we incorporated proxies of variables that might shape the wolf distribution (Table 1). Vegetation composition can indirectly influence the probability that a site becomes colonized (Marucco 2009) as well as altitude (Llaneza et al. 2012, Falcucci et al. 2013). Using the CORINE Land Cover^®^ database (U.E – SOeS, Corine Land Cover, 2006), we defined 3 covariates to characterize the landscape of the study area: forest cover, farming cover and rock cover. We used the IGN BD_ALTI^®^ database (250m resolution) to calculate the mean altitude of each site as well as the proportion of altitude higher than 2500m. Above this limit, most of the vegetation cover is grassland or rocky area. Altitude may be linked to colonization. We also predicted a site with a high proportion of high-altitude (>2500m high) would be less attractive for the species. Forest cover may structure the ungulate distribution (i.e. prey species). As a consequence, we expected that a site with higher forest cover would have a higher probability of being colonized and a site with higher rock cover would have a lower probability of being colonized. We also used the proportion of agricultural area as a covariate combining all types of farming activities including pastures areas. Those areas can be occupied by sheep, a possible prey to wolves, and therefore may have a positive influence on the settlement of the species at a site. Altitude may be linked to colonization. We also predicted a site with a high proportion of high-altitude (>2500m high) would be less attractive for the species.

**Table 1:** Description and expected effects of covariates used to describe the occupancy dynamics of wolf in France.

| Covariate | Abbreviation | Parameter | Description | Expected effect | Reference |
| --- | --- | --- | --- | --- | --- |
| Forest cover | Forest | Colonisation ($\gamma$) | Percentage of mixt, coniferous or deciduous forests cover | + | Oakleaf *et al.* 2006,<br>Fechter & Storch 2013 |
| Farmland cover | Agr | Colonisation ($\gamma$) | Percentage of pasture lands and other farming activities cover | +/- | Glenz *et al.* 2001 |
| Rock cover | Rock | Colonisation ($\gamma$) | Percentage of rock cover | - | |
| High altitude | Halt | Colonisation ($\gamma$) | Proportion of altitude higher than 2500 meters | - | Glenz *et al.* 2001 |
| Altitude | Alt | Colonisation ($\gamma$) | Mean altitude | +/- | Llanneza *et al.* 2012<br>Falcucci *et al.* 2013 |
| Distance to the closest barrier | Dbarr | Colonisation ($\gamma$) | Minimal distance between a highway or one of the five main rivers in France. | - | Falcucci *et al.* 2013 |
| Short distance occupied neighboring cells | SDAC | Colonisation ($\gamma$) | Proportion of observed occupied contiguous cells | + | Bled *et al.* 2011 |
| Long distance | LDAC | Colonisation ($\gamma$) | Proportion of observed occupied cells within a 150 | | |
| occupied |  |  | km radius without the contiguous cells. | + |  |
| neighboring cells |  |  |  |  |  |
| Year | Trend-year | Extinction $(\varepsilon)$ | Year as a linear effect | - | Marucco, 2009 |
| (continuous) |  |  |  |  |  |
| Sampling effort | SEff | Detection $(p)$ | Number of observers per site per year | + | |
| Road density | Rdens | Detection $(p)$ | Percentage of site covered by roads | + | |
| Month-survey | survey | Detection $(p)$ | Occasion of survey (categorical) | +/- | Marucco, 2009 |

Dispersal capacity is a key factor to explain the dynamic of wolf colonization (Boyd and Pletscher 1999, Kojola et al. 2006, Ciucci et al. 2009). Because cells occupied by established packs may act as a source of dispersers, (Yackulic et al. 2012), the neighborhood of an unoccupied cell may influence its colonization probability (Veran et al. 2015). On the other hand, wolves’ strategies in colonization aim at avoiding neighbors via long-distance dispersal to avoid territorial competition with neighboring packs. In that spirit, the presence of individuals at short and long-distance could be accounted for by using conditional autoregressive models and auto-logistic models (Bled et al. 2013). However, due to the computational burden and convergence issues, we could not implement this approach here. We therefore defined two covariates that consisted of the observed number of contiguous observed occupied cells at both short and long-distances around the focal cell. The short-distance covariate was defined as the number of observed occupied cells that were directly contiguous to the focal cell i.e., situated within a distance of 10 km. The limit for the long-distance parameter was set to avoid a dilution effect due to the small number of observed occupied cells at very long-distances but large enough to account for most long-distance observed occupied cells that could play a role in the colonization probability. Based on observations of wolf dispersal in the Western Italian Alps (Marucco and McIntire 2010), we set this limit at 150 km around the focal cell. We expected a positive effect of these two covariates on the probability of a site to be colonized.

Because dispersal could be driven by the presence of physical barriers (Wabakken et al. 2001, Blanco et al. 2005), we defined a landscape covariate depicting the distance from the center of the site to the closest barrier defined as highways or rivers (U.E – SOeS, Corine Land Cover, 2006). We expected this covariate to impact colonization negatively.

In the first few years after sites become newly colonized, extinction probability is expected to be high as long as only isolated individuals use them. Once a pack has settled, pack persistence is the rule for wolves when other packs are present in the surrounding areas (Mech and Boitani 2010). Pack splitting may rise from various sources including harvest or poaching of alpha pairs (Gehring et al. 2003, Brainerd et al. 2008) leading to a locally extinct site. Within the distribution of an actively expanding population, extinct sites might be recovered by surrounding individuals, either by dispersers or by neighboring packs. We therefore expected the extinction probability to decrease over time, which was tested by using “year” as a continuous covariate.

Finally, in addition to sampling effort, we considered the potential effect of road densities on the species detectability, first through facilitation of site accessibility for the observers and second, because cross roads are often used as marking sites (Barja et al. 2004), which can lead to an increase in the species detection probability. Because presence signs rely partly on track records in the snow, we considered an ‘occasion effect’ to account for the variation in detection conditions due to weather variations across the survey months (Marucco 2009).

Last, we considered the initial occupancy probability as constant since only two sites were occupied in the first year of the study, which was not enough to assess the effects of covariates on this parameter.

### Model fitting, selection and validation

We performed covariate selection using stochastic search variable selection (SSVS; George and McCulloch 1993, O’Hara and Sillanpää 2009). In brief, SSVS builds a model that includes all covariate combinations as special cases. In practice, this is achieved by adding binary indicator variables, α_p_ equals 1 or 0, which allows the estimation of the regression parameter β_p_ or excludes it by setting it to a constant (Supplementary material Table C1). In a Bayesian framework, we excluded a regression parameter by constraining it to 0 by specifying an informative prior centered on 0, while we estimated it by using a flat prior, that is β_p_ ~ (l — α_p_)Normal(0,0.0001) + α_p_Uniform0,1) with α_p_ ~ Bernoulli(0.5). Prior to model selection, we ran a Spearman test to check for correlations among covariates.

**Table C1:**
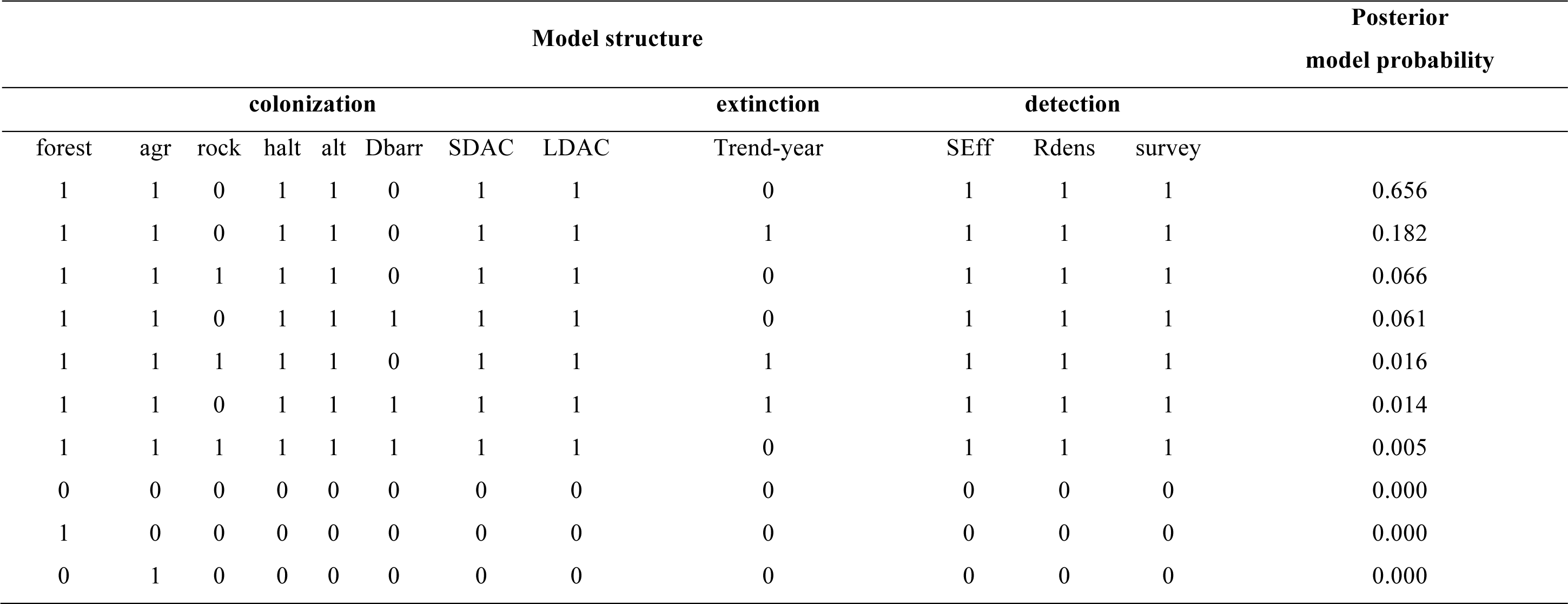
Top 10 models incorporating habitat covariates for the wolf detection/non-detection data. In the model structure, a 1/0 indicates the presence/absence of the corresponding covariate in the colonization, extinction of detection probability. Note that the intercept is always included in the model and therefore not represented in this notation.

We used the software JAGS (Plummer 2003) and Markov chain Monte Carlo (MCMC) simulations for model selection and parameter estimation. We ran three MCMC chains with a burn-in period of 2500 iterations followed by 10000 iterations on which we based our inference. We checked convergence visually by inspecting the chains and by checking that the R-hat statistic was below 1.2 (Gelman and Shirley 2011). We finally produced distribution maps of the latent states by using *a posteriori* means of the *z_i,k_* from the best model. To assess the fit of our final model, we used the posterior predictive checking approach (Gelman et al. 1996) that has recently been applied to occupancy models (Broms et al. 2016b) (Supplementary material Fig. B1).

**Figure B1:**
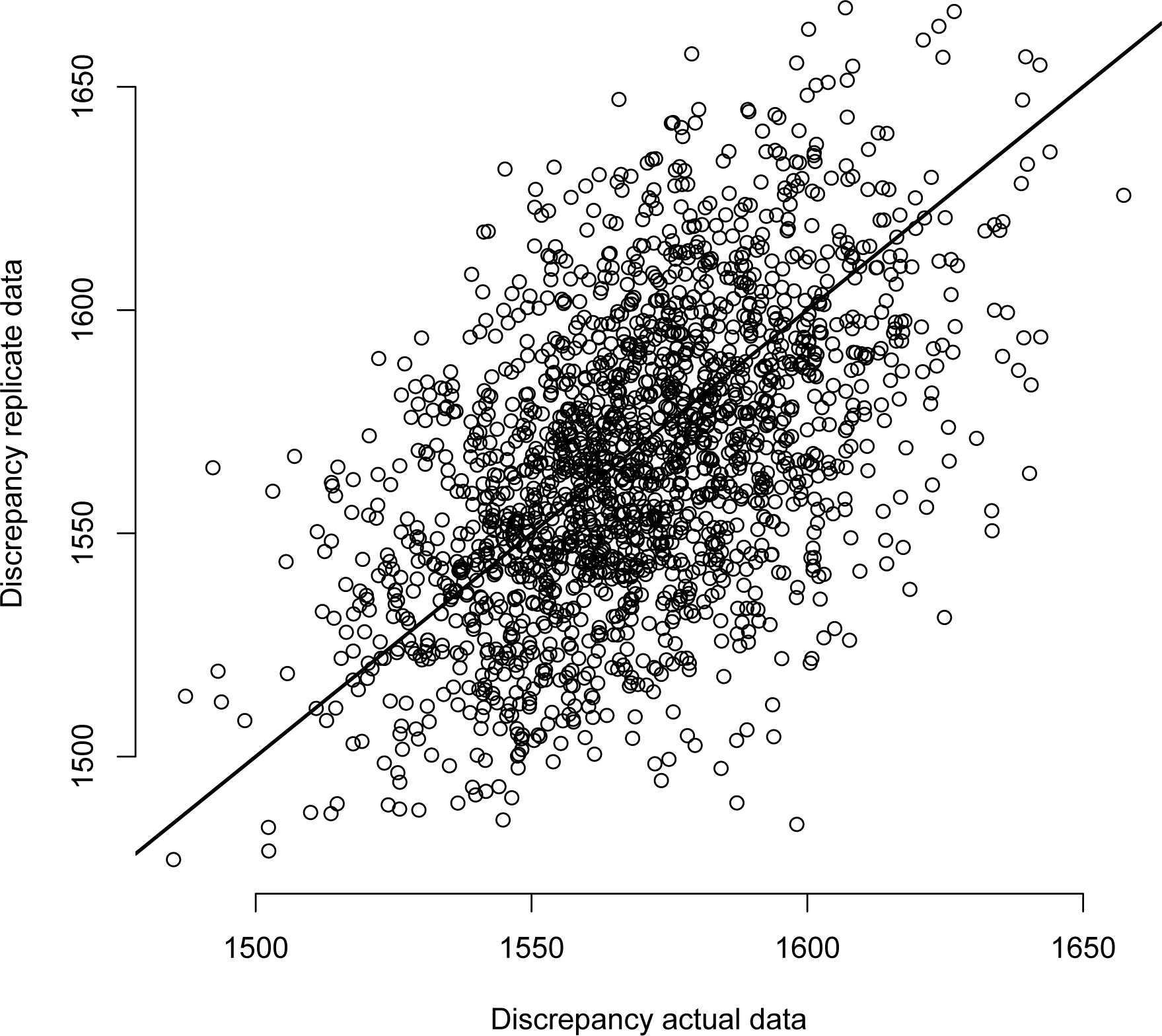
Results from posterior predictive checks for the dynamic occupancy model best supported by the wolf data. We show a scatterplot of the predicted chi-square discrepancy between simulated and expected data (on the Y axis) versus the observed chi-square discrepancy between expected and observed data (on the X axis) across MCMC samples. The Bayesian predictive p-value is 0.46 and represents the proportion of samples above the diagonal. Overall, the fit of the model seems satisfactory.

## Results

### The effect of covariates on detectability and the dynamic of occupancy

The model best supported by the data had detection as a function of sampling effort, road density and occasion (month) and colonization as a function of forest cover, farmland cover, mean altitude, proportion of high-altitude and the number of observed occupied cells at a short and long-distance neighborhood (Supplementary material, Table C1). This model appeared to fit the data adequately well (Supplementary material, Fig. B1). Posterior medians and 95% credible intervals are given for each parameter. To calculate the effect of a covariate, we set the other covariates to their mean value.

Initial occupancy probability was low, as expected since few sites were detected as occupied at the beginning of the study (Supplementary material, Table C2).

**Table C2:**
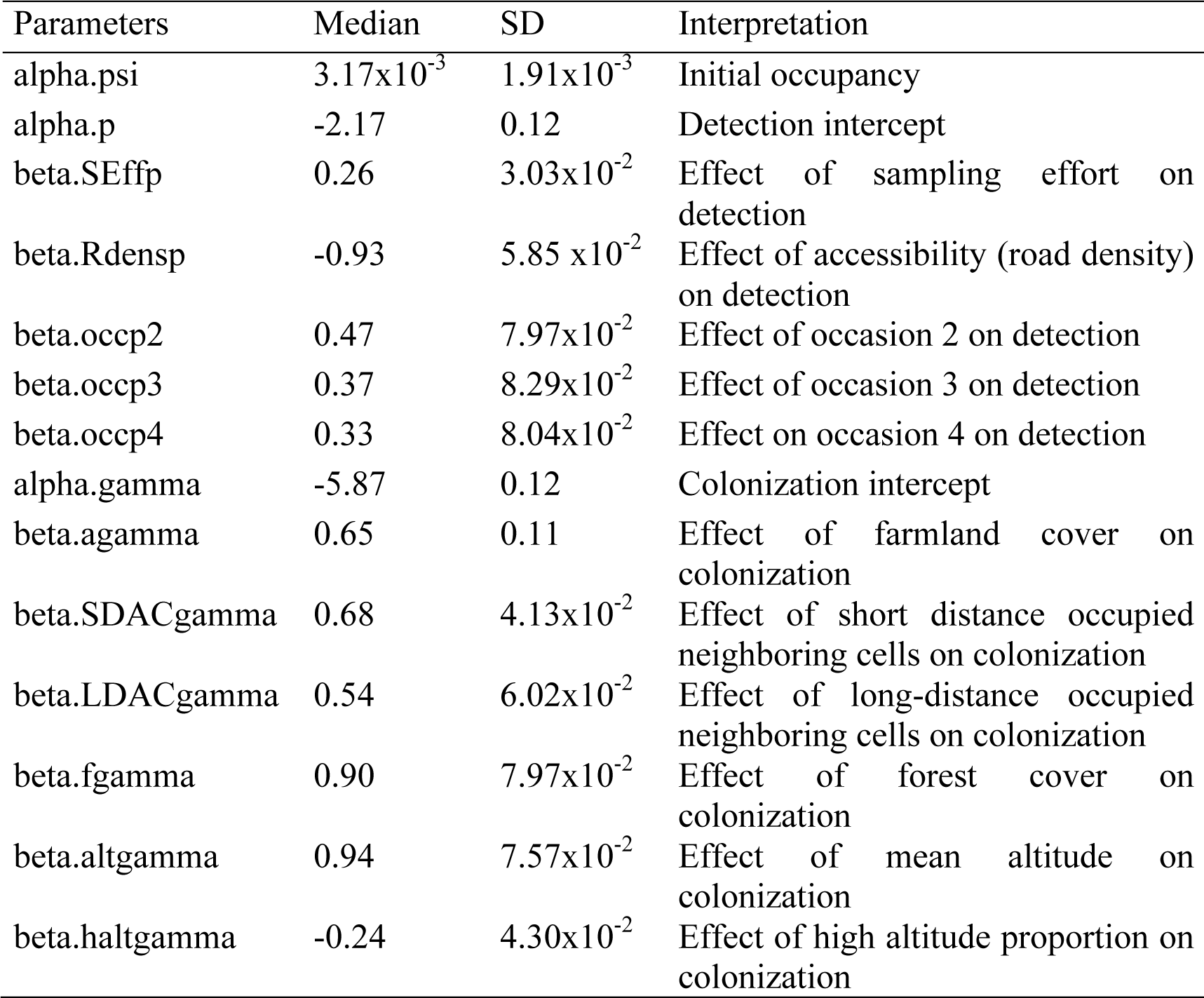
Parameters estimates from the best dynamic site-occupancy model for wolf in France between 1993 and 2014. The median a posteriori is given with the associated standard deviation (SD). Occasions 2, 3 and 4 correspond to January, February and March. Estimates are given on a logit scale except for alpha.psi which is given on its natural scale, i.e. [0, 1].

As predicted, forest cover had a positive influence on the probability that a site became colonized. The proportion of farmland area within a cell also appeared to have a positive but weak influence on this probability. Below 1500m of altitude, the probability that a site became colonized was close to zero, whereas above this limit the probability reached up to 0.26 (0.16; 0.41) (Fig. 2). This probability decreased with the high-altitude proportion in a site. Finally, as predicted, both the short and long-distance count of observed occupied neighboring cells had a strong influence on the probability that a site became colonized over a year and was dependent of the early vs. late period of the wolf recovery trend. Over time, the number of observed occupied neighboring cells increased at both short and long-distance (Supplementary material, Fig. D1). If all of the 8 neighboring cells were observed as occupied, the probability that the target site became colonized was 0.37 (0.23; 0.54) compared to a colonization probability of 2.71×10^−3^ (2.11×10-3; 3.47×10-3) if the target site had only 0 to 2 contiguous neighboring cells observed occupied. As this number increased, the probability that a site became colonized increased accordingly (Fig. 2).

**Figure 2:**
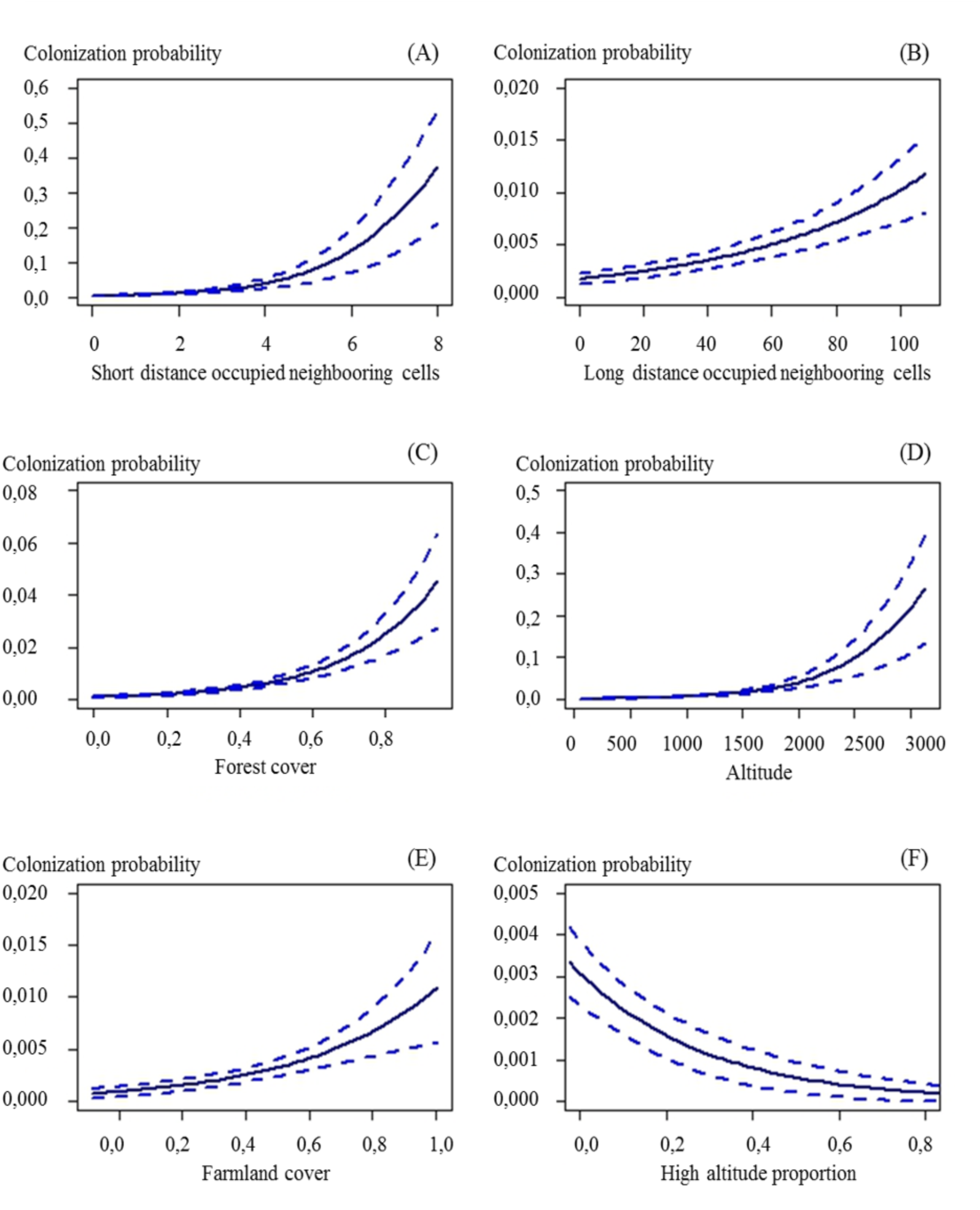
Relationship between the estimated colonization probability and (A) short-distance occupied neighboring cells, (B) long-distance occupied neighboring cells, (C) forest cover, (D) altitude, (E) farmland cover, and (F) site proportion of altitude higher than 2500 m.

**Figure D1:**
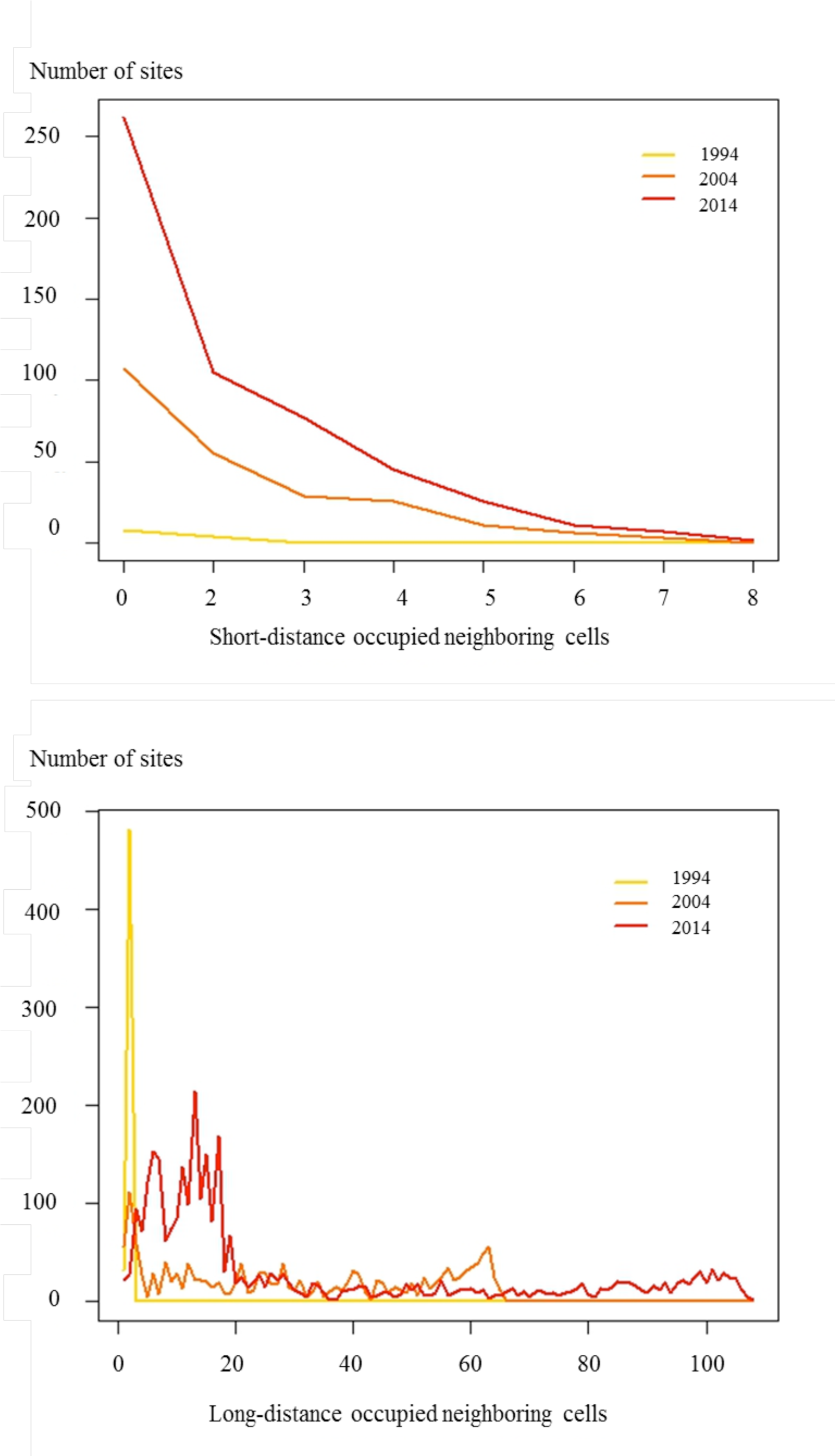
Number of sites having more than 0 observed occupied neighboring cells at short (contiguous cells) and long distance (between 10 km and 150 km) for 1994, 2004, and 2014.

Sites located within the Alps had the highest number of observed occupied sites at both short and long-distance. Colonization probability was the highest in this area (Fig.3). The highest part of the Alps (i.e. sites with the greatest proportions of high-altitude) remained with a low colonization probability (Supplementary material, Fig. D2). Overall, this probability was higher than zero in mountainous areas and increased with time as the number of occupied sites increased (Fig. 3).

**Figure 3:**
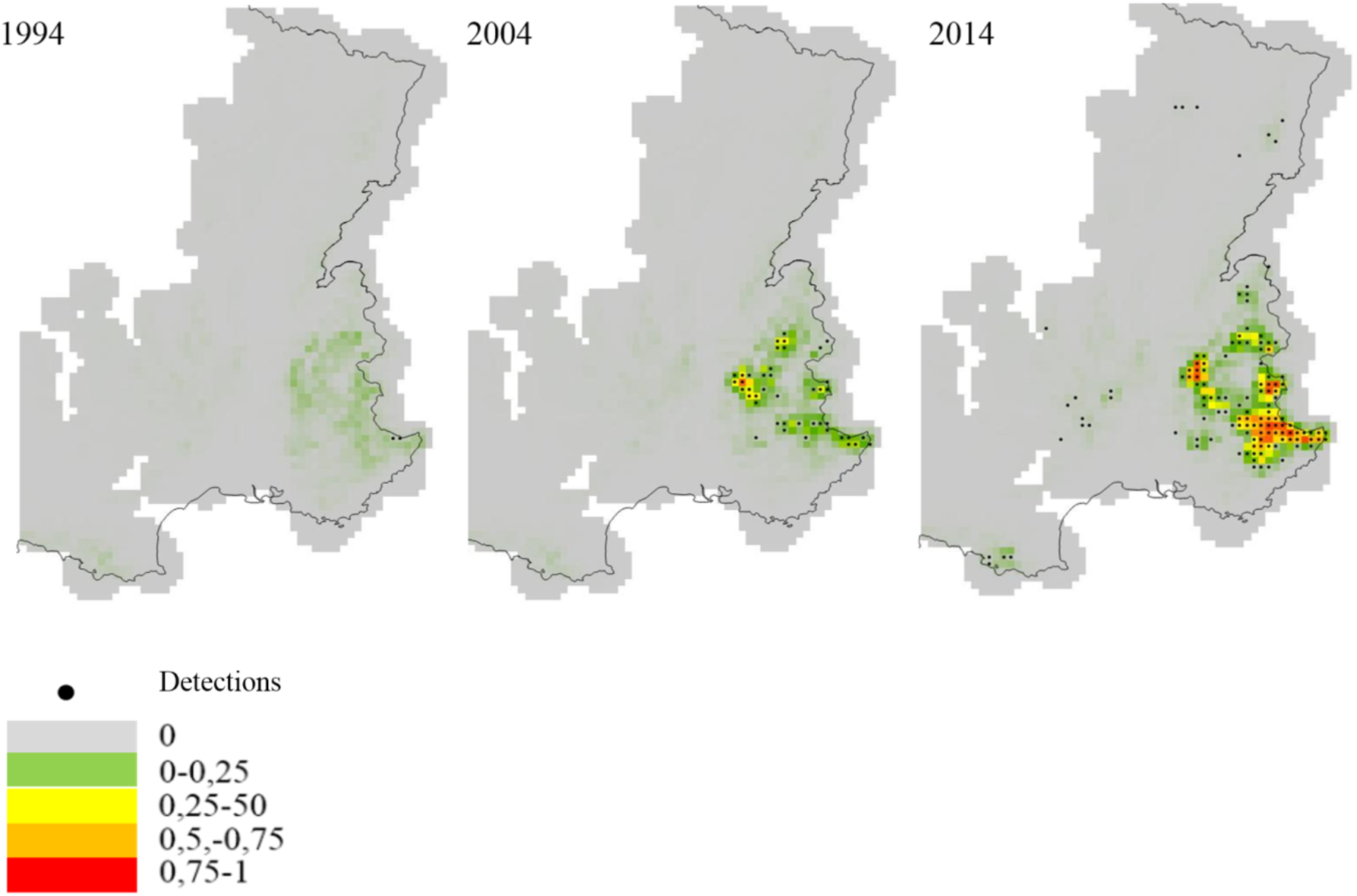
Maps of estimated colonization probability between 1993 and 1994, 2003 and 2004 and 2013 and 2014 from the best model (Table 2). Black dots represent detections made in 1993, 2003 and 2013.

**Figure D2:**
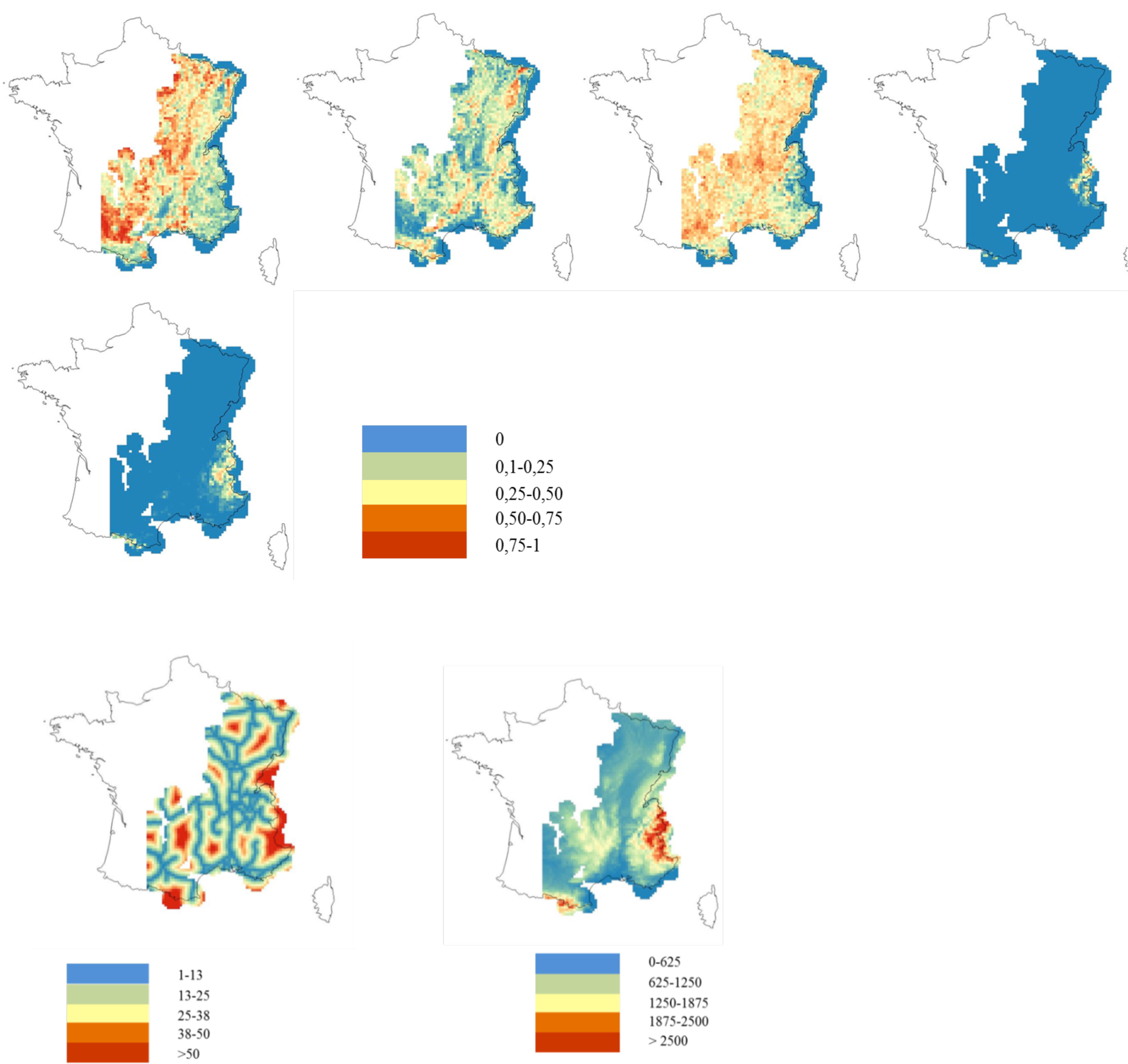
Maps of covariates tested in the study. First row from left to right: proportion of farmland cover (from 0 to 1); proportion of forest cover (from0 to 1); road density (from 0 to 1); high altitude density (from 0 to 1). Second row: proportion of rock cover (from 0 to 1). Third row: left: distance from the cell center to the closest barrier (highway or river); right: mean altitude (in meters).

Finally, and as expected, detection probability varied according to the survey month with the lowest mean value of 0.07 (0.06; 0.09) in December and the highest value of 0.11 (0.09; 0.14) in January, and intermediate values of 0.10 (0.08; 0.13) in February and 0.10 (0.08; 0.12) in March when road density and sampling effort were set to their mean values (Fig. 4). As expected, detection probability increased when the number of observers per site increased but, in contrast with what we expected, decreased with increasing road density. The sensitivity analysis showed weak effects of variations in the prospection areas used to build the sampling effort, except for the number of observed occupied sites (Supplementary Material, Fig. A3).

**Figure 4:**
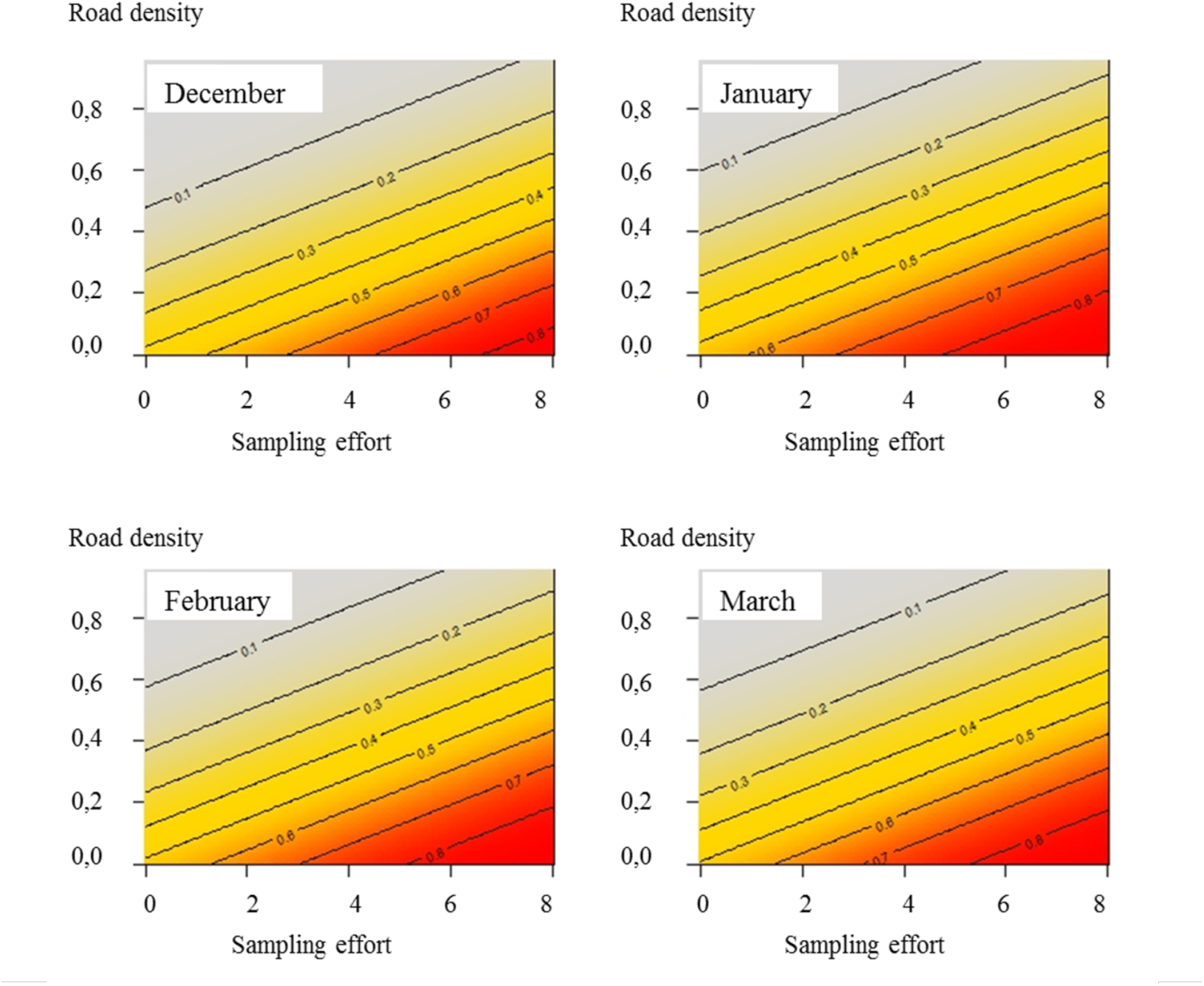
Joint effects of road density, sampling effort and occasion (month) on the species detection probability.

### Distribution map

From 1992 to 2014, 13,554 presence signs were recorded by the network and used in our analysis. The species was initially spotted in 2 cells in 1993 and was detected in 143 cells in 2014 (around 70-fold increase, see top panel in Fig. 5). This led to an apparent occupancy (proportion of occupied sites on the total number of sites in the study area) varying from 0.001 in 1993 to 0.046 in 2014.

**Figure 5:**
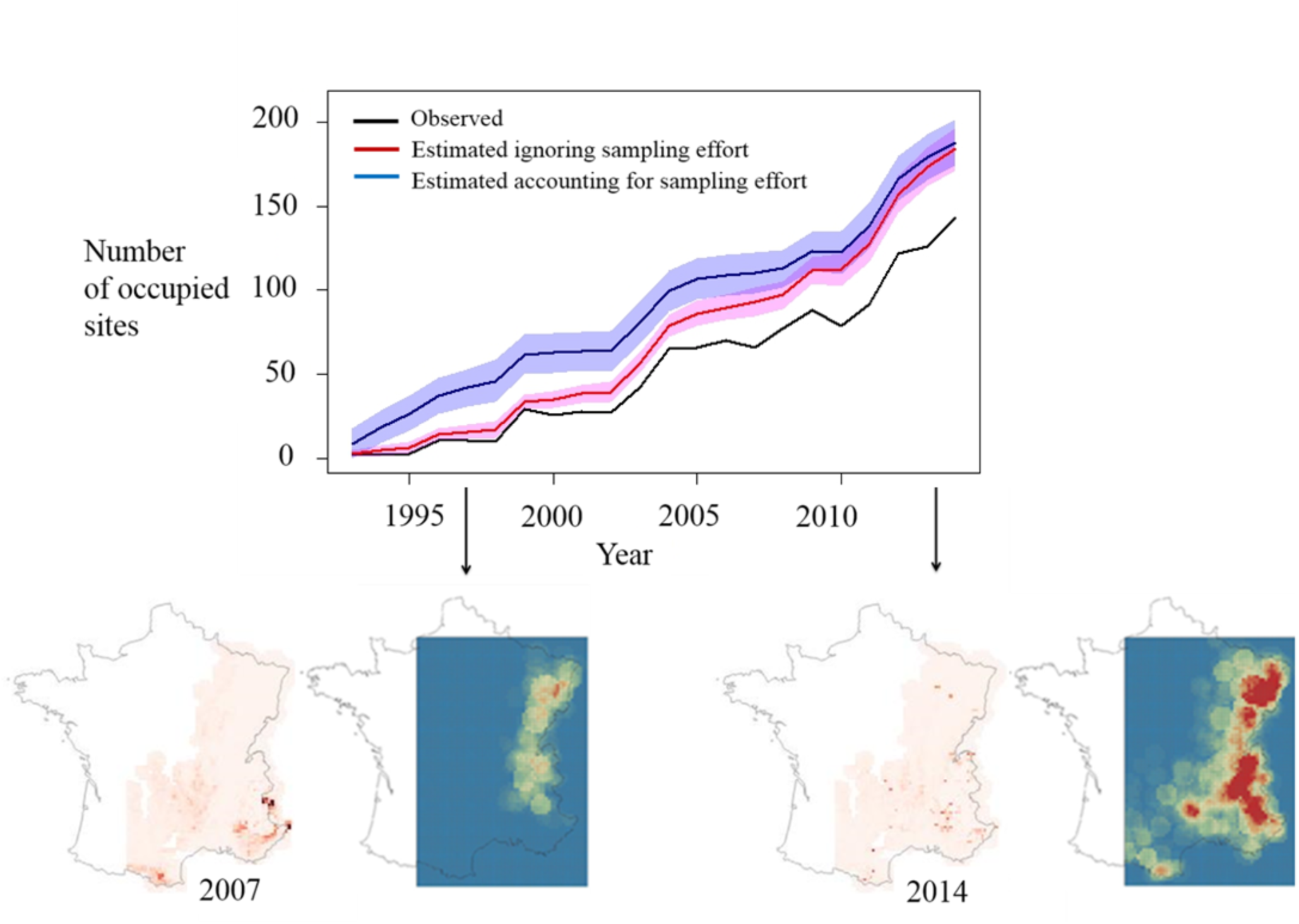
Up: Number of 10 × 10 km quadrats observed (black), estimated occupied ignoring sampling effort (red) and estimated occupied accounting for sampling effort (blue). Blue and Pink parts represent the 95% credible interval associated to the estimated number of occupied sites. Down: Maps of differences between estimates of occupancy from the model accounting for sampling effort and the one ingnoring sampling effort. Dark red sites are sites that appeared estimated occupied by the model accounting for sampling effort but did not appear occupied once ignoring sampling effort. Both maps are associated with maps of the sampling effortfor the years 1997 and 2014.

Accounting for both sampling effort and imperfect detection, we estimated the number of occupied sites as up to 10 (1; 20) in 1993 and up to 193 (178; 208) in 2014 (top panel in Fig. 5); overall, the estimates were higher than the naïve estimates of occupancy. When we ignored the sampling effort in the detection process, we found an estimated number of occupied sites equal to 3 (1; 5) in 1993 and up to 184 (171; 196) in 2014 leading to an average of 9 (8; 10) newly occupied sites per year. Most discrepancies between the two models (accounting for vs. ignoring the sampling effort) were found at the early stage of the colonization process when the network of observers was implemented mainly in the southeastern part of the Alps (compare bottom left and right panels in Fig. 5; see also Supplementary material, Fig. D3). Accounting for the sampling effort allowed us to infer the species presence on sites that were not prospected or prospected with a low sampling effort. As soon as the number of observers increased, the network was more homogeneously spread in space and estimates of the number of occupied sites became similar whether the sampling effort was included in the model or not (top panel in Fig. 5).

**Figure D3:**
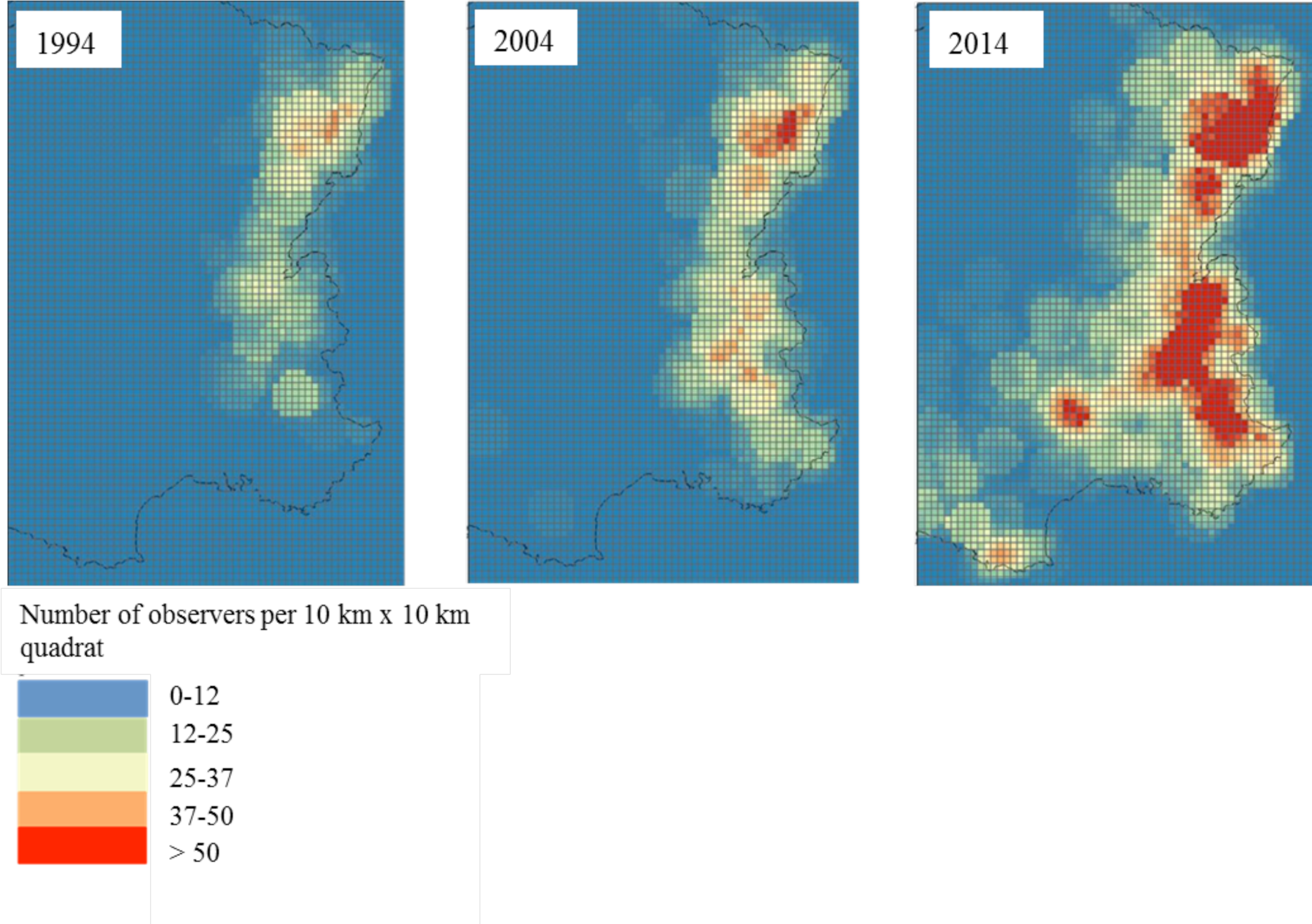
Maps of sampling effort for 1994, 2004, and 2014. Sampling effort was defined as the number of prospecting observers per site per year. In 1994, 1036 sites were prospected by at least one observer and 3083 were prospected in 2014. In 1994 only sites in North-Eastern part of France were prospected and in 2014 all Eastern France was prospected with some parts in South-West. Size of the prospection area depended on the socio-professional category of observers: observers from an administrative field (policemen for instance) were assigned a theoretical prospection area the size of the French department in which they were affiliated, observers from national parks, regional natural parks and natural reserves were assigned a prospection area the size of those areas. Observers from the farming profession and hunters were assigned a prospection area the size of the county they work in. Scientists, members of naturalist associations and private observers were assigned an area a quarter the surface of the French department where they found signs. Observers working in the French Game and Wildlife Agency (ONCFS) were assigned an area half the surface of the French department they work in. Observers from the French National Office for Forest (ONF) were assigned an area 1/10 the surface of the department they work in.

Our results showed that in 1994 the species was found only in the Southern Alps, then actively colonized towards the Northern Alps at the beginning of the 2000’s. The colonization process started to reach the Pyrenees and Massif Central area in early 2000 and the Vosges area in the very northern part of France, at the beginning of the 2010’s,indicating that the French wolf population is still in a phase of expansion. On average 9 (6; 12) new sites became occupied per year with a minimum of 0 newly occupied sites in 2002 and a maximum of 29 newly occupied sites in 2012. This led to an average growth rate (i.e. number of new sites divided by the total number of sites the previous year) of 18% (15%; 19%) (Fig. 6). This growth rate decreased over time, from 125% at the early stage of the wolf colonization in 1994 to 5% in 2014, but the species is still in an expanding phase mainly thanks to the colonization outside of the alpine range.

**Figure 6:**
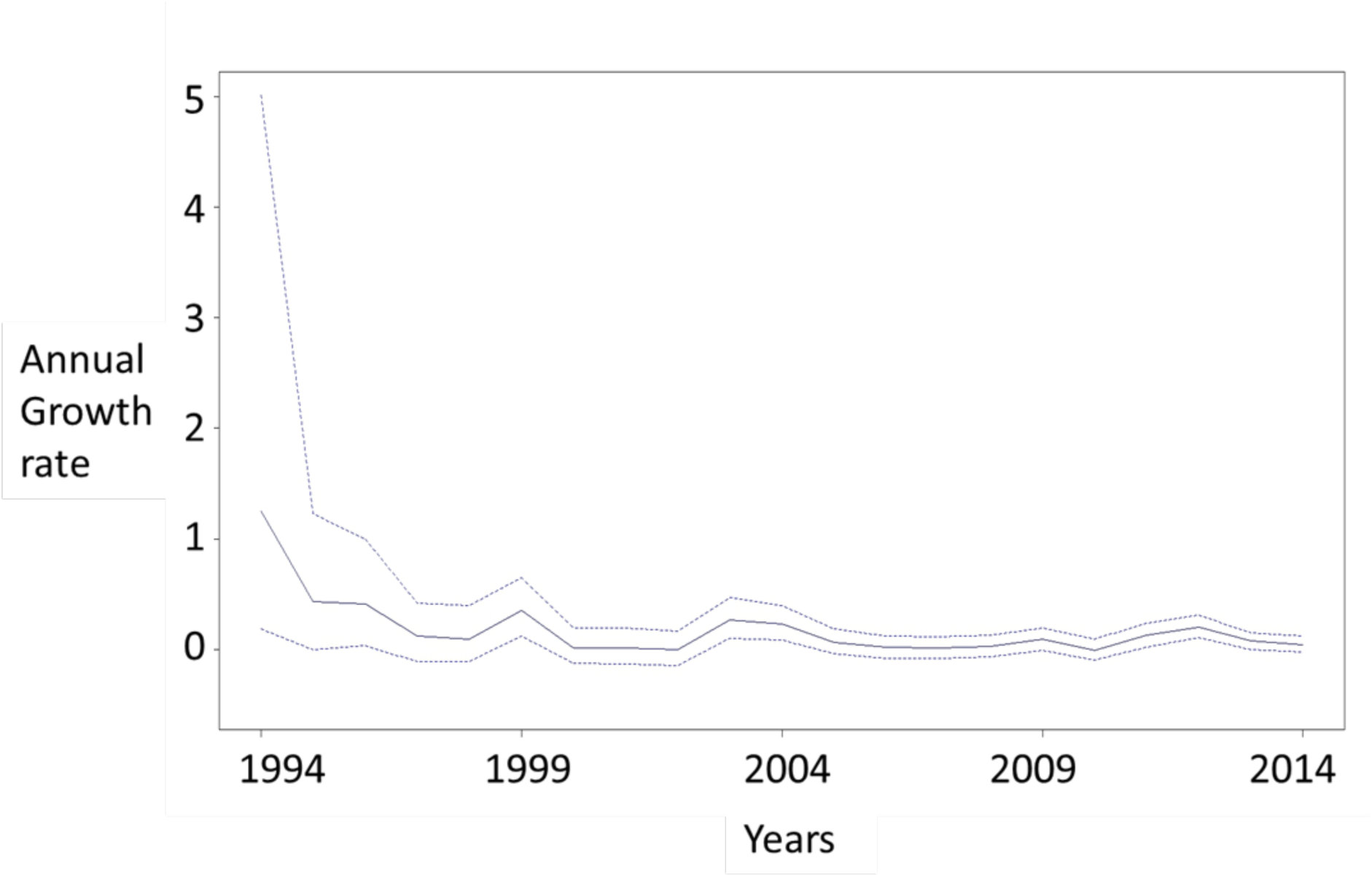
Growth rate (i.e. number of new sites rated by the total number of sites the previous year, multiplied by 100) given for each year from 1994 to 2014.

The model did not predict absence in places where presence signs were found (Fig. 7). Sites with high occupancy probability were mainly those close to the sites where the species had been previously detected, mostly due to the effect of short-distance neighbors. Some sites had a high probability of being occupied (> 0.75), however the uncertainty associated with those predictions was also high (standard deviation [SD] > 0.30). We found sites with high probability of occupancy (> 0.75) with low uncertainty (SD < 0.20), and some of those sites were observed as occupied in the following year because the model propagates information backwards in time and so *z_k_* is informed directly by *z_k_*_+1_.

**Figure 7:**
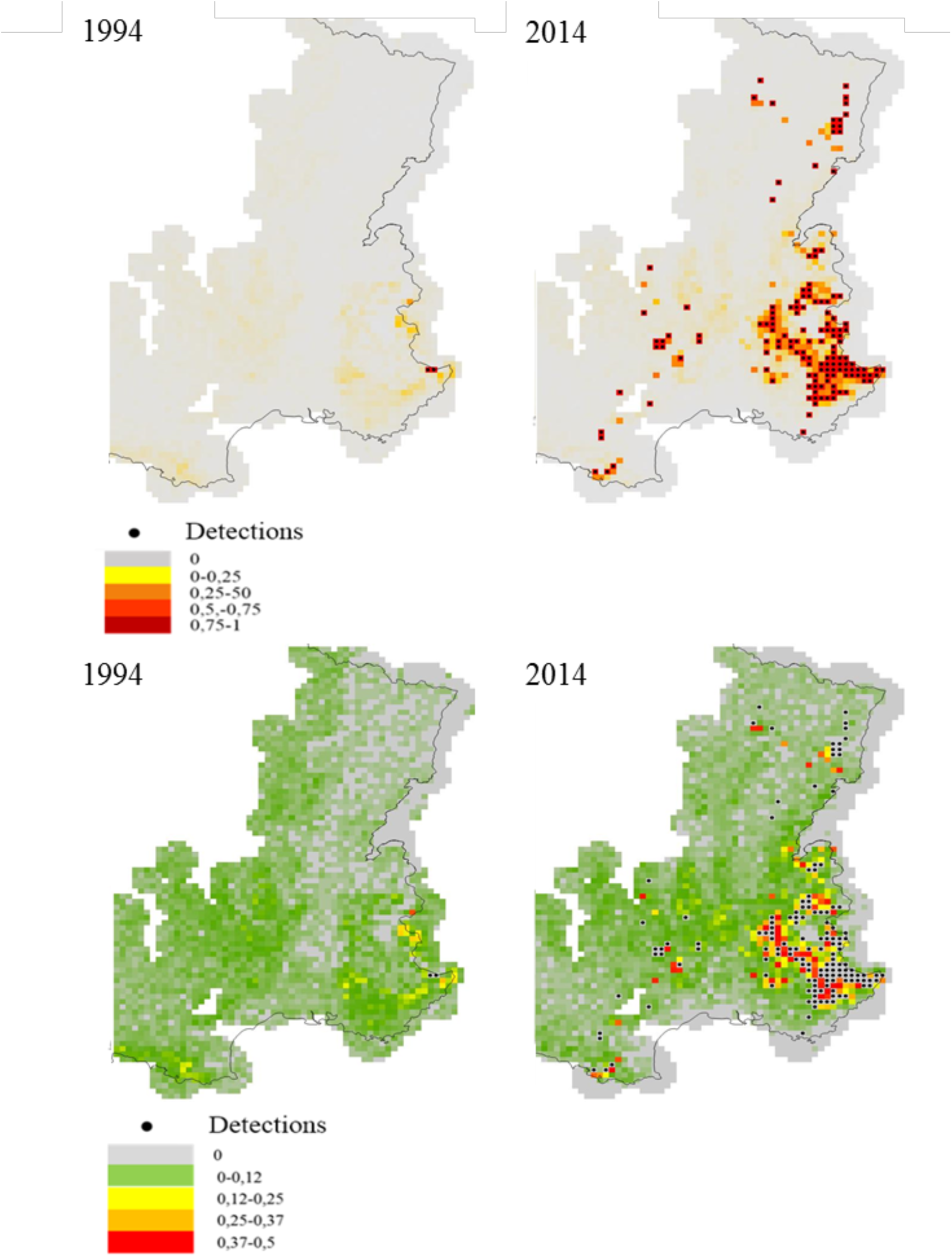
Maps of estimated occupancy (top) and associated standard deviation (bottom) for years 1994 and 2014. Black dots represent detections made in 1994 and 2014.

## Discussion

Determining favorable areas is often accomplished by building distribution maps using habitat suitability models (e.g., Mladenoff et al. 1999) or occupancy models (e.g., Marucco 2009). However, these studies often rely on a static relationship between the species of interest and its environment (Jedrzejewski et al. 2008). Here, we used dynamic site-occupancy models and brought new insights on the processes governing the dynamic of recolonization of a keystone carnivore species. By controlling for species detectability and heterogeneous sampling effort, our approach can be used to assess the distribution dynamics of any species based on opportunistic data, pending relevant information is gathered on the people collecting the data.

### Model assumptions

Site occupancy models rely on several assumptions that need to be discussed (Mackenzie et al. 2003, 2006). First, the species should not be detected when absent from a site (i.e. no false positives). This is unlikely to happen in our case since we did not account for presence signs that were rejected because they did not fulfill the standardized criteria used to avoid species misidentification (Duchamp et al. 2012). If doubts persist about the occurrence of false positives, this assumption could be relaxed by using site-occupancy models that account for misidentifications (Miller et al. 2011, Rich et al. 2013). Second, detection histories of all sampling units are assumed to be independent. However, detection histories were likely dependent in space because of a non-homogeneous spatial sampling effort inherent to opportunistic data. We partly accounted for this non-independence by quantifying the sampling effort. Furthermore, by accounting for the number of observed occupied neighboring cells, we made the detection history of a focal cell dependent partly on the detection histories of the neighboring cells. If the source of dependence is unknown, spatial autocorrelation can be modeled using geostatistical tools on occupancy or extinction/colonization parameters and also on detection (Bled et al. 2013). Third, the status of a site should not change during primary occasions - the closure assumption (Rota et al. 2009). If movements or mortality occurred inside or outside of the sampling sites, it is likely that the probability of occupancy in a given time interval did not depend on the occupancy status of a site in the previous time interval (Mackenzie and Royle 2005). In this situation of so-called random temporary emigration, the bias in parameter estimates is minimal, but occupancy should be interpreted as use of the sampling area rather than the proportion of area occupied by the species (Mackenzie et al. 2004). To prevent for this potential bias, we used the data provided within the winter period from November to March as a primary occasion because it corresponds to the most stable period in the social organization of the packs, between two main dispersal events in October (pup integration into the packs’ hunting activities) and the next mating season in March. Fourth, there should be no unmodelled heterogeneity in the model parameters. In our study, heterogeneity might remain in species detection even after accounting for spatio-temporal variation in the sampling effort or in the colonization parameter after accounting for the effect of environmental covariates. Regarding the detection probability, some heterogeneity might remain due to a difference of detection in the presence signs, e.g. tracks vs. hair (Graves et al. 2011). This was unlikely to occur in our study because the vast majority of presence signs are tracks. Regarding the colonization parameter, even though we had data on the number of killed preys during the hunting season, we did not have information on wild prey density at such a large scale. During winter (i.e. our primary occasions), wild preys consist mainly of Chamois (*Rupicapra rupicapra*), mouflons (*Ovis ammon*), roe deer (*Capreolus capreolus*) and red deer (*Cervus elaphus*) (Duchamp et al. 2012), for which we used characteristics of their habitats as a proxy for their presence (Jedrzejewski et al. 2008).

Besides the usual assumptions of occupancy models, we also had to deal with opportunistic data that are collected through non-standardized sampling protocols. To cope with opportunistic data, we defined a grid of spatial units that was overlaid on the map of detections/non-detections. In doing so, the size and shape of these sampling units might have an impact on inference about the wolf distribution. Indeed, if the size of the sampling unit is too small, then there is a risk of having very few detections within a year, which would make the estimation of the detection probability difficult. On the other hand, if the size of the sampling units is too large, then there is at least one detection in any cell, which is of little use to estimate the distribution. We used 10×10km cells as sampling units, a choice we made in agreement with what was recommended by the European Union and also shown to be the best tradeoff between the species home range and sensitivity of the distribution to the size and shape of the unit cell (Marboutin et al. 2010). The average size of wolf home ranges vary between 100 and 400 km^2^ in Western and central Europe (Ciucci 1997, Mech and Boitani 2010, Duchamp et al. 2012). Although these cells might not entirely cover wolves territories, Latham et al. (2014) studied the effect of grid size to assess wolf’s occupancy and found that taking a large grid size may not be appropriate for areas with moderate to high wolf density as it can overestimate occupancy rate.

Last, we assumed that observers were prospecting homogeneously inside the prospection area we assigned to them. This assumption may have been violated for two reasons. First, we believe that an observer was more likely to prospect more intensively near the center of the prospecting area, because it was defined as a home or work location, or near places where she/he already found presence signs. We also assumed that observers were prospecting homogeneously in time. This hypothesis can also be violated because observers may show different patterns in sampling frequency and some might not be prospecting during the months of winter. Finally, we made the assumption that once entered in the network, observers did not leave it unless we had information indicating the contrary such as a change of job or social status. Consequently, we might have overestimated the number of observers actually prospecting in the network for a given year. We therefore recommend recording carefully the activity of observers within the network to get a realistic picture of the actual sampling effort (Beirne and Lambin 2013).

### Effects of environmental covariates

We used road density as a proxy of human presence and found a negative influence on the detection probability. When defining the road density covariate, we accounted for all types of roads (except highways) and assumed that this covariate could be a proxy for site accessibility. Because many observers from the network are wildlife professionals who are familiar with opportunistic surveys, main roads may not be used and accessibility to a site may consist mostly in dirt and forest roads or pathways. The negative influence could be explained by the fact that a high road density may also affect the spatial distribution of wolves. Because wolves tend to avoid roads (Whittington et al. 2005), there might be fewer presence marks. As expected, we found that detection probability increased when sampling effort increased, therefore highlighting the importance to account for imperfect detection when it is likely to be inhomogeneous in time and space. Finally, detection varied according to the month of the survey, which can be explained by the variability in snow conditions in the study area, with higher detectability within the alpine range than in the Massif Central or lowlands for instance.

We found that colonization was mainly influenced by the number of observed occupied neighbors at short and long-distances, showing that dispersal is a key factor of the dynamic of occupancy. These results corroborate those of Adams et al. (2008) who showed that dispersal was the main component explaining wolf population dynamics. Several long-distance dispersal events have been documented across the alpine area (Wolf Alpine Group. 2014) and in France from the very south eastern part to the northern Alps, from the Alps to the Massif Central or the northern Alps to the very northeastern part of France (Duchamp et al. unpublished data). Further studies explicitly modeling dispersal processes could help to better predict wolves colonization by accounting for factors that could enhance or slow down the dispersal rate for instance (Broms et al. 2016a).

We found that mean altitude had a positive effect on colonization probability. Wolves are highly flexible and are able to live in various areas from maize cultures to high mountains (Kazcenski et al. 2013). Starting from Italy westward to the Alps (Lucchini et al. 2002, Fabbri et al. 2007), wolves reached the alpine range via the natural Apennine mountain corridor. Therefore, the effect of mean altitude may be related to the history of the wolves’ natural recovery process. However, we also found a negative effect of the proportion of altitude higher than 2500m, i.e. the higher the proportion of high-altitude, the less likely a site was to become colonized. Above 2500m, high vegetation turns from forested ecosystems to sparse vegetation, above alpine meadows with rocky covers and snow. In contrast, more forest cover associated with lower altitudes (<2500m) increased the probability that a site become colonized mainly because these habitats’ structure and composition are much more suitable to the presence of key prey species like deer (Suter et al. 2004) or mountain ungulates (Darmon et al. 2012). To a lesser extent, the effect of farmland cover was also found to have a positive but weak influence on the colonization probability, possibly because pasture areas host domestic preys during the summer period (Meriggi and Lovari 1996). The inclusion of more explicit covariates related to pastoral activity, such as the number of sheep in space, may provide a better understanding of the interaction between domestic prey and wolf presence, but these were not available to us.

### Trends in wolf recolonization

Colonization patterns have been studied during recent decades (Wabakken et al. 2001). It appears that in Scandinavia, wolves were showing a colonization process that is typical of species with high dispersal capacities and pre-saturation dispersal (Swenson et al. 1998). This process is characterized by single long leaps forward and as a consequence, the colonization front is less well defined (Hartman 1994) compared to a stepping stone dispersal strategy. Wolves seem to follow a similar pattern in France (Fig. 6) with an effect of long-distance neighborhood on colonization probability. This biological trait used by wolves is mainly known as a mechanism to avoid competition with other packs (Hayes and Harestad 2000). Once the area becomes more saturated, dispersers may fulfill remaining gaps in between other established territories, settling at long-distance unoccupied sites with higher risks of mortality due to an Allee effect (Hurford et al. 2006, Sanderson et al. 2013) or demographic stochasticity (Vucetich et al. 1997). In line with Marescot et al. (2011) who estimated a positive rate of increase in abundance, we demonstrated that the spatial dynamic mechanism of the wolves’ natural recovery is still going on, particularly outside the alpine range both northward and westward. However, this recovery appeared to slow down in proportion with the total number of occupied sites per year, mainly due to sites becoming saturated within the alpine range and/or a recent increase in official wolf controls held by the government aimed to deter damage on livestock.

We used dynamic occupancy models to assess the *current* and *dynamic* distribution of a species that is expanding since it returned; there is a temptation to go a step forward and aim at forecasting its *future* distribution. However, we emphasize the difficulty of achieving this objective because we used environmental variables to explain colonization of the species that had already occurred; by definition, we could not incorporate the drivers that may appear relevant to explain future colonization events. For instance, now that wolves have settled in the alpine range and continue to expand, they are likely to encounter new environments such as lowlands in the next few years, a landscape that may drive future colonization. Consequently, use our model as a predictive tool should be considered in an adaptive framework, i.e. by updating the management rules and the distribution maps every year during the active colonization phase.

The outcomes of our analyses have important consequences for managing animal species because their conservation statuses must be assessed partly through trends in their distributions (see art.1 of the Habitats Fauna Flora European Directive). Dynamic occupancy models are therefore relevant tools to the decision-making process by providing maps and spatio-temporal trends. In the case of the wolf, these models can help in preventing damage to livestock (Miller 2015). The identification of areas where the species may or may not occur along with the surrounding uncertainty may be used to target specific sites and determine priorities for implementing mitigation measures.

## Acknowledgements

The authors are sincerely thankful to all the volunteers belonging to the Large Carnivore Network for the data collection and local investment in the fieldwork. J. Louvrier is thankful to the GDR Ecologie Statistique, University of Montpellier and ONCFS for grants she received to conduct her work. We warmly thank Ian Renner for editing our manuscript.

## REFERENCES

Adams, L. G. et al. 2008. Population dynamics and harvest characteristics of wolves in the Central Brooks Range, Alaska. - Wildl. Monogr. 170: 1–25.

Alberto Meriggi, S. L. 1996. A Review of Wolf Predation in Southern Europe: Does the Wolf Prefer Wild Prey to Livestock? - J. Appl. Ecol. 33: 1561–1571.

Araújo, M. B. and Guisan, A. 2006. Five (or so) challenges for species distribution modelling. - J. Biogeogr. 33: 1677–1688.

Barja, I. et al. 2004. The importance of crossroads in faecal marking behaviour of the wolves (Canis lupus). - Naturwissenschaften 91: 489–492.

Beirne, C. and Lambin, X. 2013. Understanding the Determinants of Volunteer Retention Through Capture-Recapture Analysis: Answering Social Science Questions Using a Wildlife Ecology Toolkit. - Conserv. Lett. 6: 391–401.

Bischof, R. et al. 2015. Wildlife in a Politically Divided World: Insularism Inflates Estimates of Brown Bear Abundance. - Conserv. Lett. 0: 1–9.

Blanco, J. C. et al. 2005. Wolf response to two kinds of barriers in an agricultural habitat in Spain. - Can. J. Zool. 83: 312–323.

Bled, F. et al. 2011. Assessing hypotheses about site occupancy dynamics. - Ecology 92: 938–951.

Bled, F. et al. 2013. Dynamic occupancy models for analyzing species’ range dynamics across large geographic scales. - Ecol. Evol. 3: 4896–909.

Boitani, L. 2010. Wolf conservation and recovery. -In: Mech, L. D., and Boitani, L. (eds.), Wolves: Behavior, Ecology, and Conservation. University of Chicago Press, pp 317–340

Boitani L. and Ciucci P. 1993. Wolves in Italy: critical issues for their conservation. -In: Promberger, C., and Schröder, W. (eds.), Wolves in Europe: status and perspectives. Munich Wildlife Society, pp 74–90.

Brained, S. M. et al. 2008. The Effects of Breeder Loss on Wolves. - J. Wildl. Manage. 72: 89–98.

Breitenmoser, U. 1998. Large predators in the Alps: The fall and rise of man's competitor. - Biol. Conser. 83: 279–289.

Broms, K. M. et al. 2016a. Dynamic occupancy models for explicit colonization processes. - Ecology 97: 194–204.

Broms, K. M. et al. 2016b. Model selection and assessment for multi-species occupancy models. Ecology. 97: 1759–1770.

Chapron, G. et al. 2014. Recovery of large carnivores in Europe’s modern human-dominated landscapes. - Science 346: 1517–1519.

Ciucci, P. et al. 1997. Home range, activity and movements of a wolf pack in central Italy. - J. Zool. 243: 803–819.

Ciucci, P. et al. 2009. Long-Distance Dispersal of a Rescued Wolf From the Northern Apennines to the Western Alps. - J. Wildl. Manage. 73: 1300–1306.

Council of the European Commission 1992. Council directive 92/43/EEC of 21 May 1992 on the conservation of natural habitats and of wild fauna and flora. -Official Journal of the European Communities. Series L 206: 7–49.

Darmon, G. et al. 2012. Spatial distribution and habitat selection in coexisting species of mountain ungulates. - Ecography 35: 44–53.

Diane K. Boyd, D. H. P. 1999. Characteristics of Dispersal in a Colonizing Wolf Population in the Central Rocky Mountains. - J. Wildl. Manage. 63: 1094–1108.

Duchamp, C. et al. 2012. A dual frame survey to assess time- and space-related changes of the colonizing wolf population in France. - Hystrix 23: 1–12.

European Commission, 2006. Assessment, monitoring and reporting under article 17 of the habitats directive: explanatory notes and guidelines. 64p.

Fabbri, E. et al. 2007. From the Apennines to the Alps: colonization genetics of the naturally expanding Italian wolf (Canis lupus) population. - Mol. Ecol. 16: 1661–1671.

Falcucci, a. et al. 2013. Modeling the potential distribution for a range-expanding species: Wolf recolonization of the Alpine range. - Biol. Conserv. 158: 63–72.

Gehring, T. M. et al. 2003. Limits to plasticity in gray wolf, Canis lupus, pack structure: Conservation implications for recovering populations. - Can. Field-Naturalist 117: 419–423.

Gelman, A. et al. 1996. Posterior predictive assessment of model fitness via realized discrepancies (with discussion). Stat. Sinica 6:733–807.

Gelman, A. and Shirley, K. 2011. Inference from simulations and monitoring convergence. - Handb. Markov Chain Monte Carlo: 163–174.

George, E., and McCulloch, R. E. 1993. Variable selection via Gibbs sampling. JASA 88: 881–889.

Gittleman, J. L. et al. 2001. Strategies for carnivore conservation: lessons from contemporary extinctions. - Conserv. Biol. Ser. 5: 61–92.

Graves, T. A. et al. 2011. Linking landscape characteristics to local grizzly bear abundance using multiple detection methods in a hierarchical model. Anim. Conserv. 14:652–664.

Grinnell, J. 1917. The niche-relationships of the California Thrasher. - The Auk 427–433.

Guillera-Arroita, G. et al. 2015. Is my species distribution model fit for purpose? Matching data and models to applications. - Glob. Ecol. Biogeogr. 1–17.

Guisan, A. and Thuiller, W. 2005. Predicting species distribution: Offering more than simple habitat models. - Ecol. Lett. 8: 993–1009.

Hartman, G. 1994. Long-Term Population Development of a Reintroduced Beaver (Castor fiber) Population in Sweden. - Conserv. Biol. 8: 713–717.

Hayes, R. D. and Harestad, A. S. 2000. Demography of a recovering wolf population in the Yukon. - Can. J. Zool. 78: 36–48.

Hurford, A. et al. 2006. A spatially explicit model for an Allee effect: why wolves recolonize so slowly in Greater Yellowstone. - Theor. Popul. Biol. 70: 244–54.

Hutchinson, G. E. 1957. Concluding remarks. Cold Springs harbor Symposia on Quantitative Biology 22: 415–427.

Imbert, C. et al. 2016. Why do wolves eat livestock?: Factors influencing wolf diet in northern Italy. - Biol. Conserv. 195: 156–168.

Isaac, N. J. B. et al. 2014. Statistics for citizen science: extracting signals of change from noisy ecological data. - Methods Ecol. Evol. 5: 1052–1060.

Jedrzejewski, W. et al. 2008. Habitat suitability model for Polish wolves based on long-term national census. - Anim. Conserv. 11: 377–390.

Jeschke, J. M. and Strayer, D. L. 2006. Determinants of vertebrate invasion success in Europe and North America. - Glob. Chang. Biol. 12: 1608–1619.

Kaczensky, P. et al. 2013. Status, management and distribution of large carnivores–bear, lynx, wolf & wolverine–in Europe. - Report to the EU Commission.

Kéry, M. 2011. Towards the modelling of true species distributions. - J. Biogeogr. 38: 617–618.

Kéry, M. and Schaub, M. 2011. Bayesian Population Analysis using WinBUGS - a hierarchical perspective.

Kéry, M. et al. 2010. Predicting species distributions from checklist data using site-occupancy models. - J. Biogeogr. 37: 1851–1862.

Kojola, I. et al. 2006. Dispersal in an Expanding Wolf Population in Finland. - J. Mammal. 87: 281–286.

Lahoz-Monfort, J. J. et al. 2014. Imperfect detection impacts the performance of species distribution models. - Glob. Ecol. Biogeogr. 23: 504–515.

Latham, M. C. et al. 2014. Can occupancy-abundance models be used to monitor wolf abundance? - PLoS One 9: e102982.

Linnell, J. D. C. and Boitani, L. 2012. Building biological realism into wolf management policy: The development of the population approach in Europe. - Hystrix 23: 80–91.

Llaneza, L. et al. 2012. Insights into wolf presence in human-dominated landscapes: the relative role of food availability, humans and landscape attributes. - Divers. Distrib. 18: 459–469.

Long, R. a. et al. 2010. Predicting carnivore occurrence with noninvasive surveys and occupancy modeling. - Landsc. Ecol. 26: 327–340.

Lucchini, V. et al. 2002. Noninvasive molecular tracking of colonizing wolf (Canis lupus) packs in the western Italian Alps. - Mol. Ecol. 11: 857–868.

Mackenzie, D. I. and Royle, J. A. 2005. Designing occupancy studies: general advice and allocating survey effort. - J. Appl. Ecol. 42: 1105–1114.

Mackenzie, D. I. et al. 2003. Estimating Site Occupancy, Colonization, and Local Extinction When a Species Is Detected Imperfectly. - Ecology 84: 2200–2207.

Mackenzie, D. I. et al. 2004. Investigating species co-occurrence patterns when species are detected imperfectly. - J. Anim. Ecol. 73: 546–555.

MacKenzie, D. I. et al. 2006. Occupancy estimation and modelling. Inferring patterns and dynamics of species occurrence. - Academic, Burlington.

Marboutin, E. et al. 2010. On the effects of grid size and shape when mapping the distribution range of a recolonising wolf (Canis lupus) population. - Eur. J. Wildl. Res. 57: 457–465.

Marcelli, M. and Fusillo, R. 2012. Land use drivers of species re-expansion: inferring colonization dynamics in Eurasian otters. - Divers. Distrib. 18: 1001–1012.

Marescot, L. et al. 2011. Capture-recapture population growth rate as a robust tool against detection heterogeneity for population management. - Ecol. Appl. 21: 2898–2907.

Marucco, F. 2009. Spatial population dynamics of a recolonizing wolf population in the Western Alps. -University of Montana, Missoula.

Marucco, F. and McIntire, E. J. B. 2010. Predicting spatio-temporal recolonization of large carnivore populations and livestock depredation risk: wolves in the Italian Alps. - J. Appl. Ecol. 47: 789–798.

Mech, L. D. and Boitani, L. 2010. Wolves: Behavior, Ecology, and Conservation: Behavior, Ecology, and Conservation. -University of Chicago Press.

Miller, J. R. B. 2015. Mapping attack hotspots to mitigate human--carnivore conflict: approaches and applications of spatial predation risk modeling. - Biodivers. Conserv. 24: 2887–2911.

Miller, D. A. W. et al. 2011. Improving occupancy estimation when two types of observational error occur: non-detection and species misidentificatio. - Ecology 92: 1422–1428.

Miller, D. A. W. et al. 2013. Determining Occurrence Dynamics when False Positives Occur: Estimating the Range Dynamics of Wolves from Public Survey Data. - PLoS One 8: e65808.

Mladenoff, D. J. et al. 1999. Predicting gray wolf landscape recolonization: logistic regression vs. new field data. - Ecol. Appl. 9: 37–44.

Molinari-Jobin, a. et al. 2012. Monitoring in the presence of species misidentification: the case of the Eurasian lynx in the Alps. - Anim. Conserv. 15: 266–273.

O’Hara, R. B., and Sillanpää, M. J. 2009. A review of Bayesian variable selection methods: what, how and which. Bayesian Anal. 4: 85–118.

Packer, C. et al. 2013. Conserving large carnivores: Dollars and fence. - Ecol. Lett. 16: 635–641.

Phillips, S. J. et al. 2006. Maximum entropy modeling of species geographic distributions. -Ecol. Model. . 190: 231–259.

Plummer, M. 2003. JAGS: A program for analysis of bayesian graphical models using gibbs sampling. - Proc. 3rd Int. Work. Distrib. Stat. Comput. (DSC 2003). March: 20–22.

Poulle, M.-L. et al. 1997. Significance of ungulates in the diet of recently settled wolves in the Mercantour mountains (southeastern France). - Rev. d’Ecologie (Terre vie) 52: 357–358.

Promberger, C., and Schröder, W. 1993. Wolves in Europe: status and perspectives. -Munich Wildlife Society.

Rich, L. N. et al. 2013. Estimating occupancy and predicting numbers of gray wolf packs in Montana using hunter surveys. - J. Wildl. Manage. 77: 1280–1289.

Ripple, W. J. et al. 2014. Trophic cascades from wolves to grizzly bears in Yellowstone. - J. Anim. Ecol. 83: 223–33.

Rota, C. T. et al. 2009. Occupancy estimation and the closure assumption. - J. Appl. Ecol. 46: 1173–1181.

Royle, J. A. and Kéry, M. 2007. A Bayesian state-space formulation of dynamic occupancy models. - Ecology 88: 1813–1823.

Sanderson, C. E. et al. 2013. With Allee effects, life for the social carnivore is complicated. - Popul. Ecol. 56: 417–425.

Schmeller, D. S. et al. 2009. Advantages of Volunteer-Based Biodiversity Monitoring in Europe. - Conserv. Biol. 23: 307–316.

Sunarto, S. et al. 2012. Tigers need cover: multi-scale occupancy study of the big cat in Sumatran forest and plantation landscapes. - PLoS One 7: e30859.

Suter, W. et al. 2004. Spatial variation of summer diet of red deer Cervus elaphus in the eastern Swiss Alps. - Wildlife Biol. 10: 43–50.

Swenson, J. et al. 1998. Geographic expansion of an increasing brown bear population, evidence for presaturation dispersal. - J. Appl. Ecol. 67: 708–715.

Thorn, M. et al. 2011. Brown hyaenas on roads: Estimating carnivore occupancy and abundance using spatially auto-correlated sign survey replicates. - Biol. Conserv. 144: 1799–1807.

Valière, N. et al. 2003. Long-distance wolf recolonization of France and Switzerland inferred from non-invasive genetic sampling over a period of 10 years. - Anim. Conserv. 6: 83–92.

Van Strien, A. J. et al. 2013. Opportunistic citizen science data of animal species produce reliable estimates of distribution trends if analysed with occupancy models. - J. Appl. Ecol. 50: 1450–1458.

Veran, S. et al. 2015. Modeling spatial expansion of invasive alien species: relative contributions of environmental and anthropogenic factors to the spreading of the harlequin ladybird in France. - Ecography 38: 001–011.

Vucetich, J. A. et al. 1997. Effects of Social Structure and Prey Dynamics on Extinction Risk in Gray Wolves. - Conserv. Biol. 11: 957–965.

Wabakken, P. et al. 2001. The recovery, distribution, and population dynamics of wolves on the Scandinavian peninsula, 1978-1998. - Can. J. Zool. 79: 710–725.

Whittington, J. et al. 2005. Spatial responses of wolves to roads and trails in mountain valleys. - Ecol. Appl. 15: 543–553.

Wolf Alpine Group 2014. Wolf population status in the Alps: Pack Distribution and trends up to 2012. -7th « Wolf Alpine Group » Workshop“wolf monitoring over the Alps – towards a unique approach” 2013 March 19- 20th - Jausiers, FRANCE.

Yackulic, C. B. et al. 2012. Neighborhood and habitat effects on vital rates: expansion of the Barred Owl in the Oregon Coast Ranges. - Ecology 93: 1953–1966.

Yackulic, C. B. et al. 2013. Presence-only modelling using MAXENT: when can we trust the inferences ? - Methods Ecol. Evol. 4: 236–243.

Yackulic, C. B. et al. 2015. To predict the niche, model colonization and extinction. Ecology, 96: 16–23.

Zurell, D. et al. 2009. Static species distribution models in dynamically changing systems: How good can predictions really be? - Ecography 32: 733–744.

